# Orthogonal proteomic platforms and their implications for the stable classification of high-grade serous ovarian cancer subtypes

**DOI:** 10.1101/793026

**Authors:** Stefani N. Thomas, Betty Friedrich, Michael Schnaubelt, Daniel W. Chan, Hui Zhang, Ruedi Aebersold

## Abstract

The National Cancer Institute (NCI) Clinical Proteomic Tumor Analysis Consortium (CPTAC) has established a two-dimensional liquid chromatography-tandem mass spectrometry (2DLC-MS/MS) workflow using isobaric tagging to compare protein abundance across samples. The workflow has been used for large-scale clinical proteomic studies with deep proteomic coverage within and outside of CPTAC. SWATH-MS, an instance of data-independent acquisition (DIA) proteomic methods, was recently developed as an alternate proteomic approach. In this study, we analyzed remaining aliquots of peptides using SWATH-MS from the original retrospective TCGA samples generated for the CPTAC ovarian cancer proteogenomic study (Zhang et al., 2016). The SWATH-MS results indicated that both methods confidently identified differentially expressed proteins in enriched pathways associated with the robust Mesenchymal subtype of high-grade serous ovarian cancer (HGSOC) and the homologous recombination deficient tumors also present in the original study. The results demonstrated that SWATH/DIA-MS presents a promising complementary or orthogonal alternative to the CPTAC harmonized proteomic method, with the advantages of simpler, faster, and cheaper workflows, as well as lower sample consumption. However, the SWATH/DIA-MS workflow resulted in shallower proteome coverage. Overall, we concluded that both analytical methods are suitable to characterize clinical samples such as in the high-grade serous ovarian cancer study, providing proteomic workflow alternatives for cancer researchers depending on the specific goals and context of the studies.

## Introduction

Advances in sample preparation workflows, mass spectrometry instrumentation, and data processing software have positioned proteomics to provide comprehensive insights into complex biological processes at a level close to the underlying biochemical mechanisms. Indeed, it is currently possible to routinely quantify >10,000 proteins in human cell proteomes (Beck et al., 2011; Nagaraj et al., 2011) and human tissue proteomes (Kim et al., 2014; Mertins et al., 2018; Wilhelm et al., 2014) using mass spectrometry-based platforms. The workflows for many of these large-scale proteomic studies entail extensive offline fractionation of the peptides generated from enzymatically-digested proteins, followed by LC-MS/MS. Consequently, proteomic analysis of large cohorts of clinical specimens (>100) requires several months for data acquisition using the afore-mentioned workflows. Further, because each fraction analyzed typically requires 1-5 microgram of total peptides, the required quantity of the original tissue sample is in the milligram level. Although more rapid proteomic workflows have been developed (Anagnostopoulos et al., 2017; Hebert et al., 2014; Kulak et al., 2014; Richards et al., 2015), they have not yet been deployed for large-scale clinical proteomic studies.

Examples of large-scale clinical proteomic studies using the 2DLC-MS/MS workflow described above include the NCI CPTAC studies. CPTAC was formed to accelerate the understanding of the molecular basis of cancer through the application of large-scale proteogenomic analyses. Several hundred tumor tissue specimens from breast, ovarian, and colorectal cancer tissues previously analyzed by NCI’s The Cancer Genome Atlas (TCGA) have also been characterized using proteomics, informed by genomics, resulting in the identification and quantification of proteins and phosphoproteins in cancer-associated cell signaling pathways and networks (Mertins et al., 2016; Zhang et al., 2014a; Zhang et al., 2016a). These studies employed data-dependent acquisition (DDA) mass spectrometry, a mode of MS/MS data collection wherein a fixed number of precursor ions whose *m/z* values were recorded in a survey scan are selected for fragmentation using a pre-determined set of rules (Mann et al., 2001). DDA-based proteomic workflows have undergone considerable optimization to improve the reliability and reproducibility of the generated data in an effort to minimize the limitations due to the stochastic nature of precursor ion selection and low sampling efficiency resulting in missing values across data sets (Mertins et al., 2018; Revesz et al., 2018; Tabb et al., 2016; Zhou et al., 2017).

A relatively newer method termed data-independent acquisition (DIA) mass spectrometry has been gaining traction in large-scale proteomic studies (Vidova and Spacil, 2017). DIA mass spectrometry is an alternative to DDA which allows all ions within a selected mass range to be concurrently fragmented and analyzed by tandem mass spectrometry. Sequential Window Acquisition of All Theoretical Mass Spectra (SWATH-MS) is an example of a DIA acquisition method whose use in proteomic studies has increased considerably within the past five years. The SWATH-MS method acquires a complete and permanent digital fragment ion record for all detectable precursor ions of a sample (Collins et al., 2013; Gillet et al., 2012; Liu et al., 2013) and introduced an iterative, targeted search strategy that determines the presence and quantity of tens of thousands of query peptides using reference fragment ion spectra for the query peptides as prior information (Rost et al., 2014). To support SWATH/DIA data analysis, several software tools have been developed and benchmarked (Navarro et al., 2016). As with any analytical methodology that has potential widespread use, several studies have been conducted to optimize and evaluate the performance of SWATH-MS (Li et al., 2017b; Rardin et al., 2015). In a multi-laboratory study including 11 sites worldwide, SWATH-MS was shown to have a linear dynamic range exceeding 4 orders of magnitude with an inter-laboratory CV of 22.0 ± 17.4% (Collins et al., 2017), demonstrating that SWATH-MS is a reproducible method for large-scale protein quantification.

To assess the potential of SWATH-MS in addressing some of the common limitations of DDA proteomic workflows, comparative analyses of SWATH-MS and DDA using isobaric tags for relative and absolute quantitation (iTRAQ) have been conducted (Basak et al., 2015; Bourassa et al., 2015; Zhang et al., 2014b). Basak *et al*. concluded that SWATH-MS and iTRAQ DDA are complementary techniques with a 60% overlap of the high confidence quantifiable proteins identified by both methods using *Saccharomyces cerevisiae* as a model system when incorporating offline peptide fractionation into the LC-MS workflow (Basak et al., 2015). In their study, Basak *et al*. incorporated first dimension separation using strong cation exchange chromatography wherein the peptides were fractionated into six fractions followed by second dimension separation using reversed-phase chromatography prior to LC-MS analysis.

In the current study, we utilized SWATH-MS to analyze peptides from 103 HGSOC tumors that were previously analyzed by iTRAQ DDA as part of the NCI CPTAC study (Zhang et al., 2016a). The iTRAQ DDA proteomic workflow resulted in the identification of 8,597 proteins from these tumors using a 24-fraction peptide separation method, whereas 2,914 proteins were quantified by SWATH-MS without peptide fractionation. We compared the two proteomic workflows on the basis of cost, robustness, complexity, ability to detect differential protein expression, and the elucidated biological information. Our analysis demonstrated that despite the greater than 2-fold difference in the analytical depth of iTRAQ DDA compared to SWATH-MS common differentially expressed proteins in enriched pathways associated with the HGSOC Mesenchymal subtype were identified by both workflows with 96% of the proteins quantified by SWATH-MS also quantified by iTRAQ DDA. We also showed that tumor subtype classification stability is sensitive to the number of samples that are analyzed. Lastly, our results indicated a conservation of the homologous recombination deficiency (HRD)-associated enriched DNA repair and chromosome organization pathways in the iTRAQ DDA and SWATH-MS data sets, thus indicating that some biological information for HGSOC could be consistently extracted from either dataset.

Taken together, compared to the DDA analysis of HGSOC using iTRAQ labeling and 2D fractionation, SWATH-MS is a robust proteomic method that can be used to re-capitulate common differentially expressed proteins in enriched pathways associated with the HGSOC Mesenchymal subtype (Zhang et al., 2016a). The SWATH-MS proteomic workflow is simpler, faster, cheaper, and consumes less sample, but it results in shallower proteome coverage. The significantly lower number of proteins detected by SWATH-MS compared to the iTRAQ DDA workflow is mitigated by the streamlined and less complex workflow, the increased sample throughput (requiring ∼80% less time), ∼10-fold reduced sample requirements and lower technical variability and attenuated signal compression. The SWATH-MS workflow therefore presents novel opportunities to enhance the efficiency of clinical proteomic studies with continuous method improvements.

## Results

### Study design and evaluation of technical performance of analytical platforms

Proteomic measurements of clinically-annotated HGSOC previously characterized by TCGA (Cancer Genome Atlas Research, 2011) were conducted using an iTRAQ DDA workflow entailing stable isotope labeling, offline fractionation, and LC-MS/MS analysis of each fraction (Zhang et al., 2016a). A total of 103 of these tumors were used for the current SWATH-MS analysis. An overview of the experimental design of our study is shown in Figure 1.

**Figure 1.**
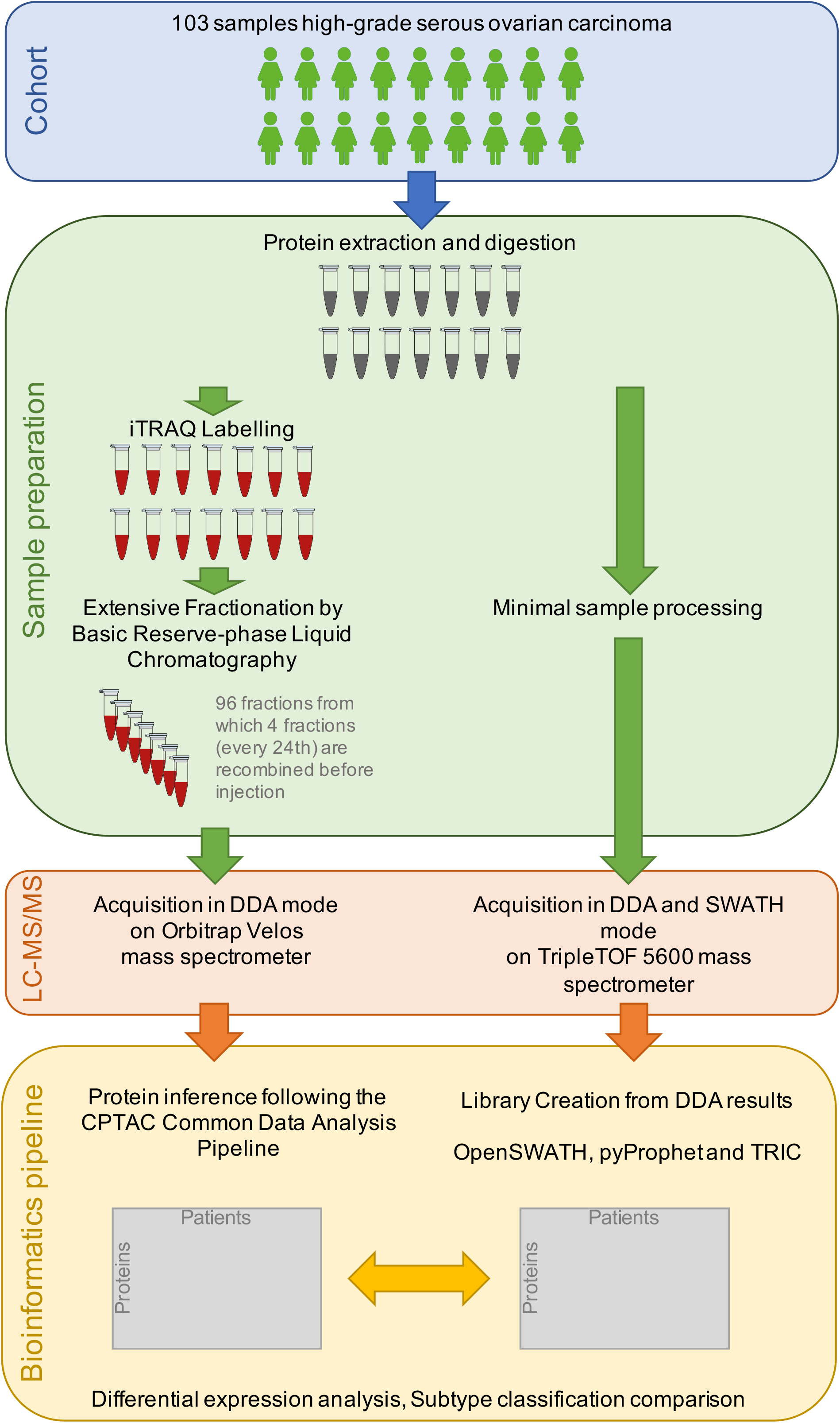
Experimental design of proteomic analysis of high-grade serous ovarian tumors using DIA mass spectrometry instrument platforms. A total of 103 clinically-annotated ovarian high-grade serous carcinomas previously characterized by The Cancer Genome Atlas (TCGA) were processed for proteomic analysis using an iTRAQ DDA method wherein the samples were subjected to fractionation prior to data acquisition and also using a SWATH-MS method without fractionation prior to analysis. The data processing and bioinformatics pipeline enabled differential expression analysis and tumor subtype classification comparison.

Protein was extracted from each tumor specimen followed by enzymatic digestion with trypsin. For the iTRAQ DDA workflow, the resulting peptides were labeled with 4-plex iTRAQ reagents followed by combination into analysis sets comprised of the peptides from three tumors, each labeled with a distinct iTRAQ tag, and an iTRAQ-labeled reference pool comprised of the peptides from most of the tumors. Each analysis set was subjected to offline fractionation into 24 concatenated fractions, and the fractions from each analytical set were sequentially analyzed using a DDA method on an LTQ-Orbitrap Velos mass spectrometer. An unfractionated aliquot of each analytical set was also analyzed by DDA-MS without fractionation. In comparison, the SWATH-MS workflow did not require stable isotope labeling. However, for the purpose of generating a spectral library to facilitate the targeted protein identification, each sample from 103 tumors was pooled, fractionated to 48 fractions and subjected to DDA analysis.

The CV of SWATH-MS data was assessed using two QC approaches (**Figure S1**). For the first method, peptides from HEK293 cells were analyzed in triplicate on three days for a total of nine runs between the DDA data acquisition runs for spectral library generation and the SWATH acquisition of the ovarian tumor data. The median CV for 3,855 quantified proteins was 8% and the mean total CV (reflecting the intra- and inter-day CV) was 15% for the nine technical repeat analyses (Collins et al., 2013; Collins et al., 2017) (**Figure S1a**). For the second QC method, peptides from a control ovarian tumor were analyzed using a DDA method on the same 5600+ TripleTOF mass spectrometer that was used to acquire the SWATH data. These QC samples were run in duplicate immediately prior to the SWATH acquisition of the ovarian tumor samples, and again in duplicate 10 days later when half of the ovarian tumor sample data acquisition runs were completed. These measurements resulted in a total CV of 7% for 781 quantified proteins (**Figure S1b**). For both QC strategies, the duration of the LC gradient was identical to the gradient used for the SWATH analysis of the HGSOC samples. Hence, the results from these QC strategies evaluated the analytical measurement/technical variability of the SWATH-MS platform that was used to acquire the data from the ovarian tumor proteins.

We compared the analytical differences between the SWATH-MS and the iTRAQ DDA workflow with respect to relative protein abundances (Figure 2A), correlation of the normalized relative abundance of the quantified proteins (Figure 2B), and variability of the constituent peptides from the quantified proteins (Figure 2C; y-axis indicates the peptide variability calculated by the standard deviation of the quantified peptides for each protein divided by the mean peptide intensity per protein). Protein abundances were determined based on the normalized log_2_ intensity ratios compared to the reference iTRAQ channel of each iTRAQ set for the iTRAQ data, and protein abundances in the SWATH-MS dataset were determined based on the log_2_ intensity of each protein as represented by the mean intensity of the constituent peptide ratios (peptide intensity divided by the mean peptide intensity) (Figure 2A). The median relative log_2_ protein abundance of the iTRAQ data was 0.01 compared to −0.23 in the SWATH-MS data. The compressed distribution of the quantified protein abundances of the iTRAQ DDA data reflects the well-documented phenomenon of ratio compression in iTRAQ-based relative quantification (Ow et al., 2011; Savitski et al., 2013) (Figure 2A).

**Figure 2.**
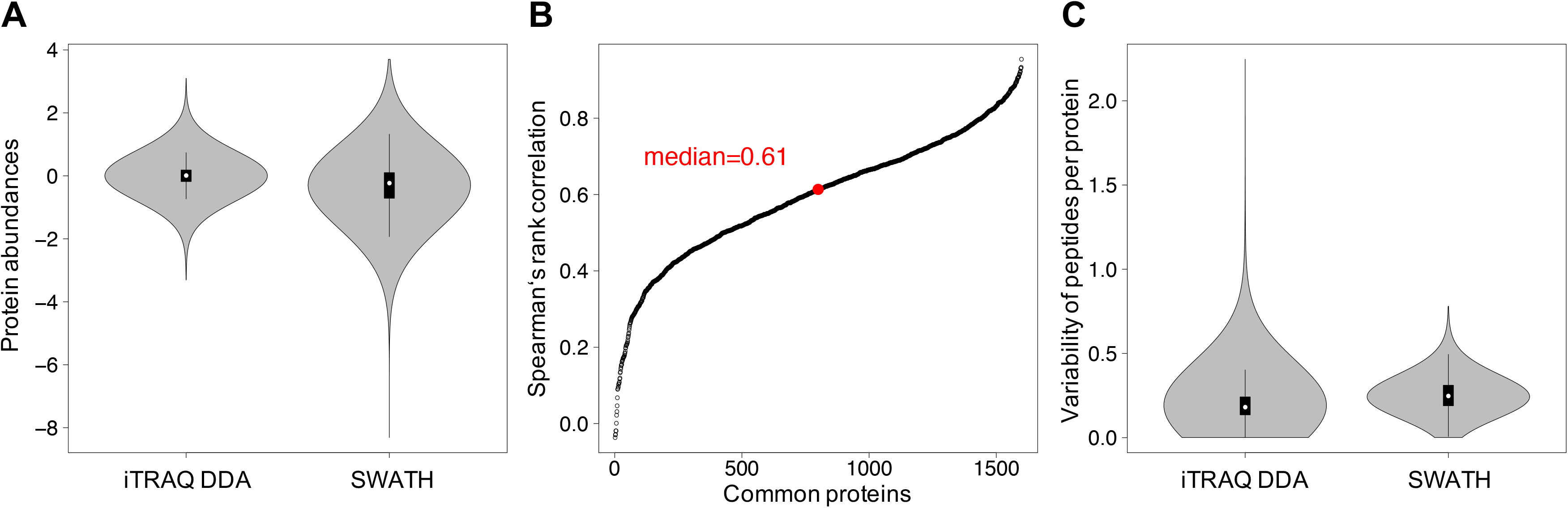
Comparison of iTRAQ DDA and SWATH-MS proteomic data based on completeness, variability and required resources. **a)** Distribution of protein abundance in the iTRAQ DDA and SWATH-MS datasets. **b)** Spearman’s rank correlation of 1,599 proteins quantified by iTRAQ DDA and SWATH-MS. **c)** Peptide variability calculated by the standard deviation of peptides per protein divided by mean peptide intensity per protein.

Spearman’s rank correlation of the relative protein abundance was used to compare the 1,599 proteins quantified by iTRAQ DDA and SWATH-MS. The median *ρ* of 0.61 indicates a moderately positive correlation (Figure 2B). Among the factors that likely preclude the median *ρ* from being higher are the fundamental differences in protein quantification (log_2_ ratio of reporter ion intensities in the iTRAQ DDA data compared to the summed fragment ion intensities and normalized to the respective mean peptide intensities for the SWATH-MS data), and the differences in protein quantification wherein multiple peptides from the same protein were used to quantify each protein in the dataset. A direct comparison of the CV of each method was not possible because replicate analyses of the samples were not conducted; therefore, a CV-like score was calculated which represents the variability of the sibling peptides from the quantified proteins. Because aliquots of identical samples were used for this analysis, the displayed variation reflects technical differences in the measurements and data handling of both methods. Although the median variability of the SWATH-MS data was slightly higher than the iTRAQ DDA data (0.25 vs. 0.18), the overall variability of the iTRAQ DDA data was much broader (Figure 2C).

Both assessed proteomic methods yielded different numbers of quantified proteins: a cumulative total of 8,597 quantified proteins resulting from the iTRAQ DDA analysis and a cumulative total of 2,914 quantified proteins resulting from the SWATH-MS analysis. A total of 1 µg of each of the 24 fractionated peptide samples was used for the iTRAQ DDA analysis, and 1 µg of peptide samples from each tumor was injected for the SWATH-MS analysis. In iTRAQ DDA, we used the original quantified proteomic data from Zhang *et al*. (Zhang *et al., 2016a)*, and we only used values obtained from unshared peptides with other proteins (proteotypic peptides). The SWATH-MS data was filtered on a peptide-query level FDR of 1% and a protein–level global FDR cutoff of 1% based on the method described by Rosenberger *et al*. for the statistical control of peptide and protein error rates in large-scale DIA analyses (Rosenberger et al., 2017a). In addition, only proteotypic peptides were used for quantification for this dataset.

For any proteomic dataset consisting of multiple analyses of biologically distinct samples, not every protein will be detected in every sample, and the resulting data matrix will have missing values. This can either be due to technical reasons, i.e., not every protein present in a sample will be identified due to the stochastic nature of the sampling of ions by the mass spectrometer, or reasons related to biological variability.

We therefore assessed the occurrence and distribution of missing values in the two data sets. **Figure S2** shows the distribution of the proteins with missing values in the iTRAQ and SWATH data. Here, we show the number of proteins with various percentages of missing values in the iTRAQ DDA (**Figure S2A**) and SWATH-MS datasets (**Figure S2B**), starting with proteins without any missing values across all 103 samples up to proteins with more than 91% missing values. In the iTRAQ dataset, approximately 50% of the proteins did not have any missing values, whereas in the SWATH-MS dataset, 11% of the proteins did not have any missing values and 50% of the proteins had ∼30% missing values. The iTRAQ DDA data set was directly filtered for the proteins without missing values (4,363). The same approach was not applicable in SWATH-MS due to the sparsity of the matrix. Only working with complete measurements in SWATH-MS would also result in neglecting potential biological effects, where proteins might not be detected due to downregulation. Due to the instrument duty cycle and the stochastic nature of data acquisition, more missing values are expected to occur in low abundant proteins than in high abundant ones (Wang et al., 2007). We assessed this as well and could observe this inverse effect, which is displayed in **Figure S2C**. Hence, we applied a filtering strategy following this assumption by allowing more missing values in high abundance proteins and less missing values among the low abundance proteins, yielding 1,659 proteins, instead of taking the 4,363 proteins without missing value from our previous iTRAQ based DDA approach. To enable the use of a complete matrix, an imputation approach was adopted based on that used in the Perseus software program (Tyanova et al., 2016). Imputation has been demonstrated to be an effective approach to address the challenge of missing values in SWATH-MS and related DIA approaches (Collins et al., 2017; Karpievitch et al., 2012; Rost et al., 2016).

The robust filtering approach used for the SWATH-MS data resulted in 1,659 proteins that were quantified with high confidence across all 103 tumors. Among these proteins, 1,599 were also quantified using iTRAQ-DDA, and these are the proteins that were used for all subsequent analyses, including the elucidation of the molecular subtype classification of the tumors based on their proteomic signatures.

### High-grade serous ovarian carcinoma subtype classification based on proteomic signatures

The rationale for the molecular subtyping of HGSOC is related to efforts to develop more specific and effective therapeutic strategies given the heterogeneity of this type of ovarian cancer. To assess the ability to classify the iTRAQ-DDA and SWATH-MS data into subtypes based on proteomic signatures, we used the same approach as reported by Zhang *et al*. wherein an unbiased molecular taxonomy of HGSOC was established using protein abundance data to identify subtypes that exhibit biological differences (Zhang et al., 2016b). The 1,599 proteins that were quantified in the iTRAQ-DDA and SWATH-MS datasets were used to classify the two datasets separately using mclust (Fraley and Raftery, 2002) based on the z-score-transformed protein abundances, and the emergent protein modules were characterized using weighted gene-correlation network analysis (WGCNA) (Langfelder and Horvath, 2008) and Reactome pathway enrichment. The heat map resulting from the clustering analysis is shown in **Figure S3A** with the proteins listed in horizontal rows and the tumors listed in vertical columns. The colored vertical bars represent the transcriptome-based HGSOC subtypes (Verhaak et al., 2013) (“Original TCGA”), the proposed proteomic iTRAQ DDA subtypes (“Original CPTAC” (Zhang et al., 2016a); “iTRAQ DDA”) and the proposed proteomic SWATH-MS subtypes (“SWATH-MS”).

The correlation of the WGCNA-derived protein modules with the proteomic subtypes derived from the iTRAQ DDA and SWATH-MS data sets is shown in **Figure S4**. The enriched Reactome pathways in the WGCNA-derived modules in the SWATH-MS data include ECM organization, immune response, metabolism, complement cascade and fibrin clot formation, and gene expression and translation. In the SWATH-MS dataset, the tumors with proteins exhibiting a significantly positive correlation (Pearson correlation coefficient *ρ* = 0.52; p-value *= 2e^-8^*) with the ECM organization Reactome pathway were assigned to the Mesenchymal subtype. A significantly positive correlation (Pearson correlation coefficient *ρ* = 0.54; p *= 5e^-9^*) was also observed among the proteins assigned to the gene expression and translation ontology.

Based on the observation highlighted by the dashed box in Figure 3 indicating a large degree of similarity among the genomic and proteomic profiles of the Mesenchymal subtype tumors, we conducted a down-sampling analysis in an effort to evaluate the stability of the Mesenchymal subtype. In this analysis, the number of tumors used for the cluster analysis was systematically decreased to determine the sample number cutoff below which the enrichment of the Reactome pathways was no longer significant. The Mesenchymal cluster was the only cluster for which the enrichment of specific Reactome pathways remained significant (p<0.01, Fisher’s exact test) when the sample number was reduced to 60% (**Figure S3B**). These results suggest that the SWATH-MS and iTRAQ DDA proteomic signatures of the Mesenchymal subtype tumors are consistent and robust.

**Figure 3.**
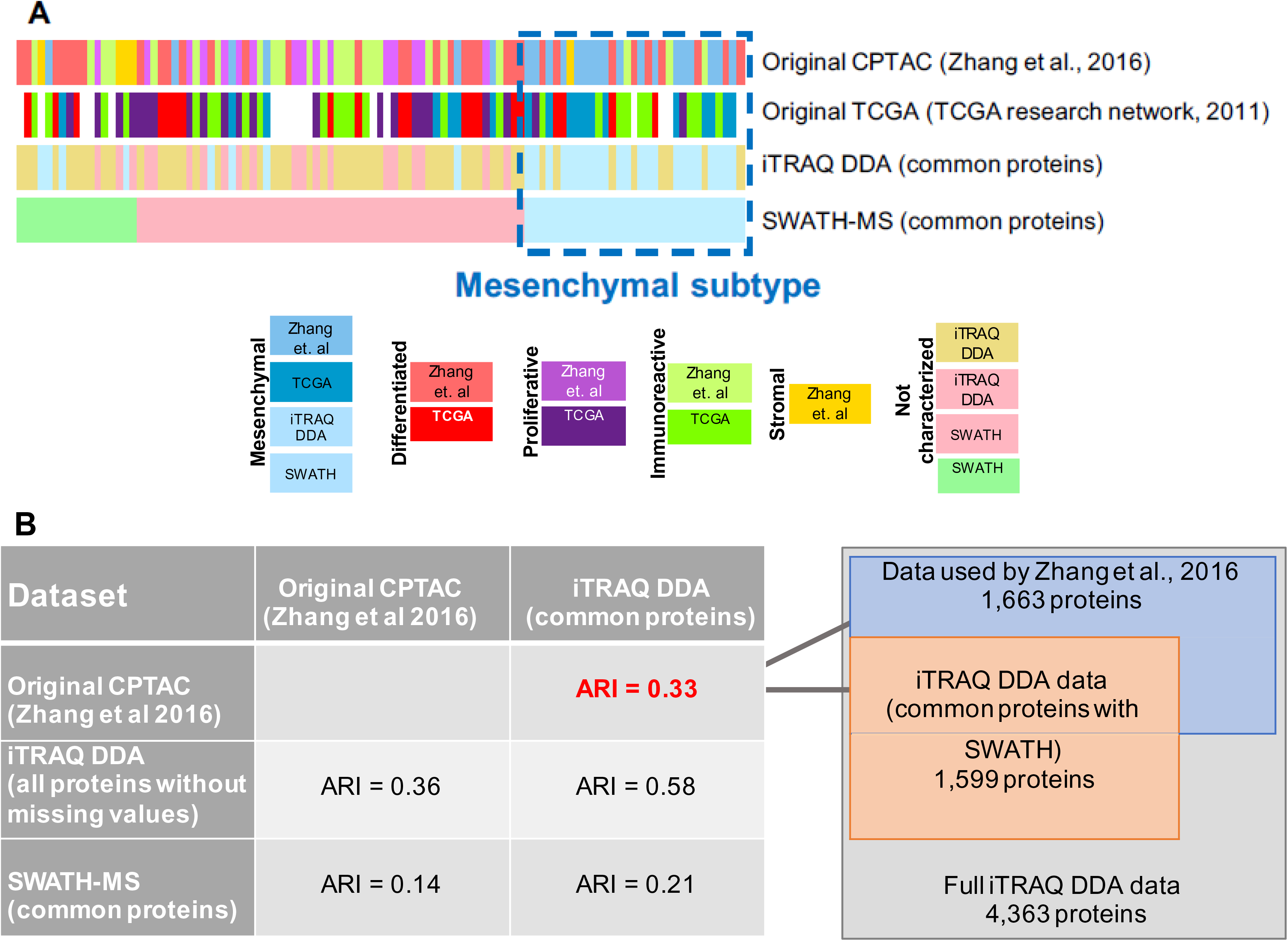
Protein (iTRAQ DDA, SWATH-MS) and mRNA-based high-grade serous ovarian cancer subtype classification. **a)** Tumor subtype classification comparison based on gene expression (TCGA study) and protein abundance (original CPTAC study (Zhang et al., 2016a), and the iTRAQ DDA and SWATH-MS data from the current analysis). Subtype classification: Blue – Mesenchymal; Red – Differentiated; Purple – Proliferative; Green – Immunoreactive; Yellow – Stromal. **b)** Subtype classification agreement among the genomic and proteomic data assessed by the Adjusted Rand Index (ARI).

Because the protein and mRNA components of the same ovarian tumors were analyzed, we were able to compare the robustness of the Mesenchymal subtype across different analytical methods. Figure 3A compares the subtyping of the tumors based on the five protein-based subtypes (Differentiated ‘D’, Immunoreactive ‘I’, Proliferative ‘P’, Mesenchymal ‘M’, and Stromal ‘S’) that emerged from the original iTRAQ DDA-based analysis of these tumors (Zhang et al., 2016a), the four mRNA-based subtypes (Differentiated, Proliferative, Mesenchymal, and Immunoreactive) that emerged from the genomic analysis (Cancer Genome Atlas Research, 2011), and the three protein-based subtypes that resulted from the clustering analysis of the iTRAQ DDA and SWATH-MS data using the 1,599 proteins that were quantified using these two analytical approaches. Based on visual analysis, the Mesenchymal subtype tumors (represented by shades of blue) exhibit the highest degree of classification agreement among the four analytical approaches. Although the accurate characterization of the molecular subtypes of HGSOC is challenging, it is widely accepted that the Mesenchymal subtype is defined by the increased expression of extracellular matrix proteins and desmoplasia (Chen et al., 2018). Compared to the other data types, the SWATH-MS data facilitates the discrete partitioning of one of the five proteomic subtypes, the Mesenchymal subtype tumors, from our previous five proteomic subtypes (Figure 3A).

### Influence of sample size and analytical depth on tumor molecular subtype classification stability

We used the Adjusted Rand Index (ARI) to quantitatively assess the agreement among the classification of the tumors based on the proteomic analyses (iTRAQ DDA and SWATH-MS) (Figure 3B). ARI values range from 0 to 1, with 1 indicating clustering results that are identical and 0 indicating clusters that are devoid of similarity. Although we did not anticipate an ARI of 1 when comparing the iTRAQ DDA and SWATH-MS datasets using the commonly quantified 1,599 proteins due to the inherent noise in biological data, we did not expect the relatively low ARI of 0.21. To further explore the similarity among the various proteomic datasets mentioned in Figure 3A, we calculated the ARI values that resulted from changing the numbers of proteins included in the analysis. The highest ARI was 0.58 which was obtained by comparing the complete iTRAQ DDA dataset without any missing values (4,363) with the 1,599 proteins in the iTRAQ DDA dataset that were also quantified by SWATH-MS. This ARI was unexpectedly low given the expected level of similarity when comparing a subset to the full set of exactly the same data. Conversely, the lowest ARI, 0.14, was obtained when comparing the full iTRAQ DDA dataset of 8,597 proteins with the 1,599 proteins that were quantified in common using SWATH-MS. This low ARI value of 0.14 is likely reflective of the fundamental differences in protein quantification between iTRAQ DDA and SWATH-MS.

We further examined the unexpected lack of similarity when comparing the subset of the iTRAQ data to the full dataset by adopting a systematic bootstrapping approach. Bootstrapping was used to randomly select a fraction of either the 103 samples analyzed by iTRAQ DDA and SWATH-MS (Figure 4A, B) or the proteins that were quantified by iTRAQ DDA (4,363) and SWATH-MS (1,659) (Figure 4C, D). ARI values based on mclust classification were used to determine the robustness of the clusters. This bootstrapping was repeated 100 times for each subset of the proteins or samples. The resulting classifications were compared to the result from the complete initial dataset, and the distribution of ARI values is shown in Figure 4 for the iTRAQ DDA (left column, A, C) and SWATH-MS datasets (right column, B, D). A lack of statistical significance among the ARI values (n.s.) indicates cluster stability, whereas cluster instability is indicated by statistically significant changes in ARI. When we bootstrapped along the dimension of sample number, the average ARI decreased dramatically with the reduced number of samples, even when comparing 90% vs. 80% of the samples in the iTRAQ DDA and SWATH-MS datasets (Figure 4A, B, respectively). However, the corresponding median ARI values of the incrementally reduced sample sizes in the SWATH-MS dataset systematically exceed the median ARI values of the iTRAQ DDA dataset (Figure 4B). These results indicate that the fidelity of molecular clusters is rather sensitive to the number of samples included in the analysis and that the clusters identified from the SWATH-MS data generally have slightly higher confidence. In comparison, when we bootstrapped along the dimension of the number of quantified proteins (Figure 4C, D) the ARI was refractory to the number of proteins included in the clustering. The difference among the ARI values did not reach statistical significance (i.e. the clusters were stable) when the number of quantified proteins was down-sampled to 30% and 50% of the initial group of proteins in the iTRAQ DDA and SWATH-MS datasets, respectively. Hence, the resulting clusters are largely insensitive to the number of included proteins.

**Figure 4.**
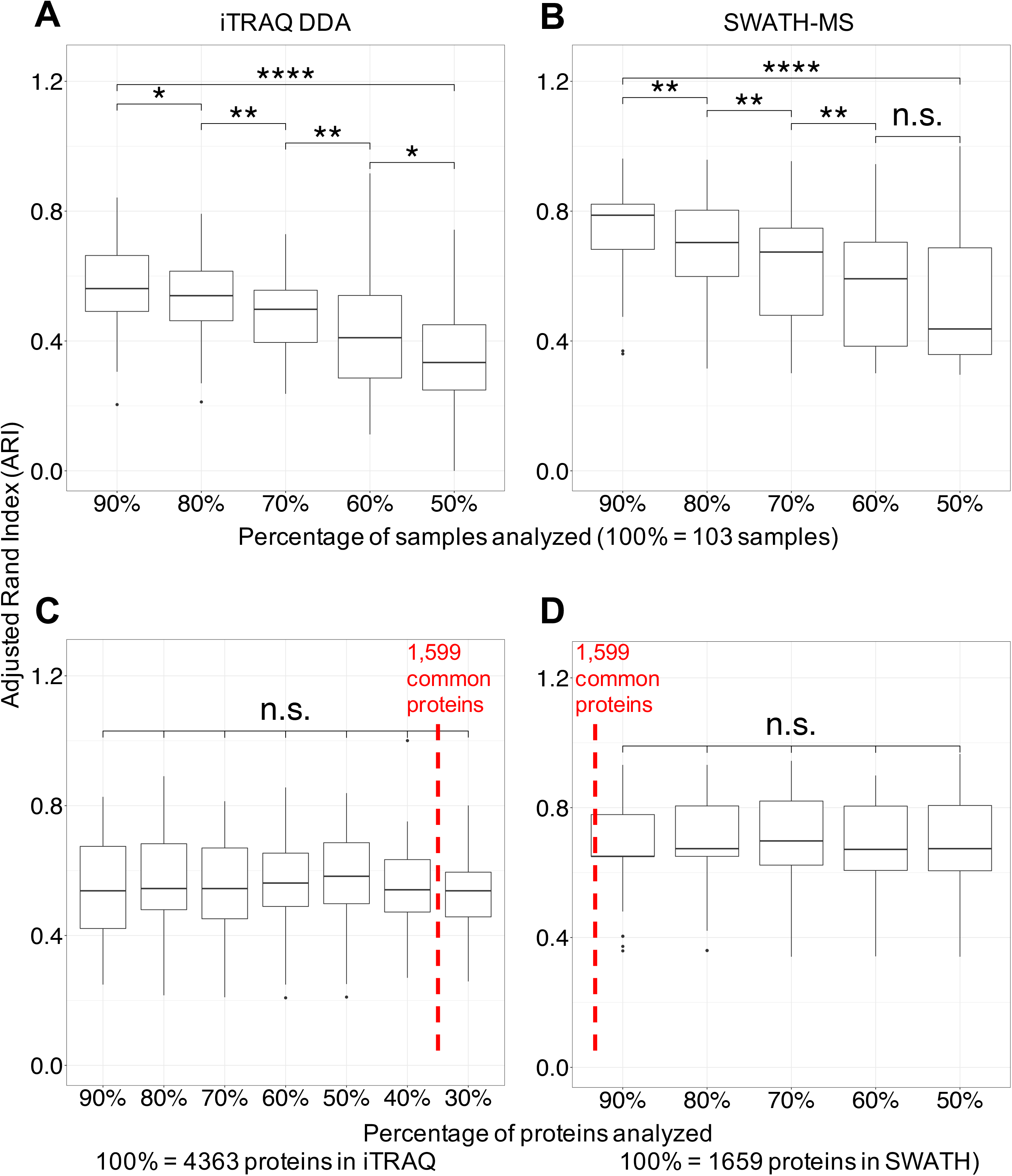
Proteomic cluster stability as a function of sample number and protein number in the iTRAQ DDA and SWATH-MS datasets. Cluster stability measured by the Adjusted Rand Index (ARI) as a function of the percentage of tumors analyzed in the iTRAQ DDA **(a)** and SWATH-MS **(b)** datasets. **c)**, **d)** Cluster stability assessed by ARI as a function of the percentage of quantified proteins in the iTRAQ DDA **(c)** and SWATH-MS **(d)** datasets. \**p*<0.05, \*\**p*<0.01, \*\*\**p*<0.001, \*\*\*\**p*<0.0001. n.s. non-significant

The sensitivity of molecular subtype clustering to the number of samples included in the analysis has been shown previously (Levine and Domany, 2001; Monti et al., 2003); however, this phenomenon has not been demonstrated in a systematic manner for proteomics data. Increasing the number of samples included in subtype classification analyses renders the classifications more robust, whereas increasing the numbers of proteins by employing proteomics methods such as 2DLC-MS/MS iTRAQ DDA does not provide an added benefit in terms of cluster stability. Our bootstrapping approach clearly demonstrated how easily classification results can be perturbed by experimental variables such as sample size, with a negative impact on the ability to extract robust biological content.

Nevertheless, in our SWATH-MS analysis, we were able to extract one robust HGSOC subtype, which was characterized by the increased relative abundance of proteins with extracellular matrix functions (**Figure S3A**). The robustness of this presumptive Mesenchymal subtype as a function of sample number was demonstrated using different subsets of samples in the SWATH-MS dataset using a Fisher’s exact test (**Figure S3B**). A fundamental premise of this stability analysis is that only the most robust information can be reproducibly extracted from the data. The emergence of the Mesenchymal phenotype as a robust subtype in the iTRAQ DDA and SWATH-MS data indicates that biologically-relevant content can be robustly extracted from these orthogonal/complementary analytical methods. The proteins characterizing the Mesenchymal cluster in the SWATH data (86 proteins) and the iTRAQ DDA data (283 proteins) were enriched in extracellular matrix organization function, show an overlap of 84 proteins (Figure 5A), and have a significantly higher correlation of the abundance of constituent proteins compared to the proteins in the entire dataset of 1,599 proteins (Figure 5B, median *ρ* = 0.79 compared to Figure 2B, median *ρ* = 0.61). The relatively high correlation of the abundance of proteins comprising the Mesenchymal cluster is also evidenced by the overlap among the Mesenchymal subtype tumors (Figure 3A). These results suggest that the robustness of the clusters increases with increasing quantification accuracy rather than by increasing number of quantified proteins.

**Figure 5.**
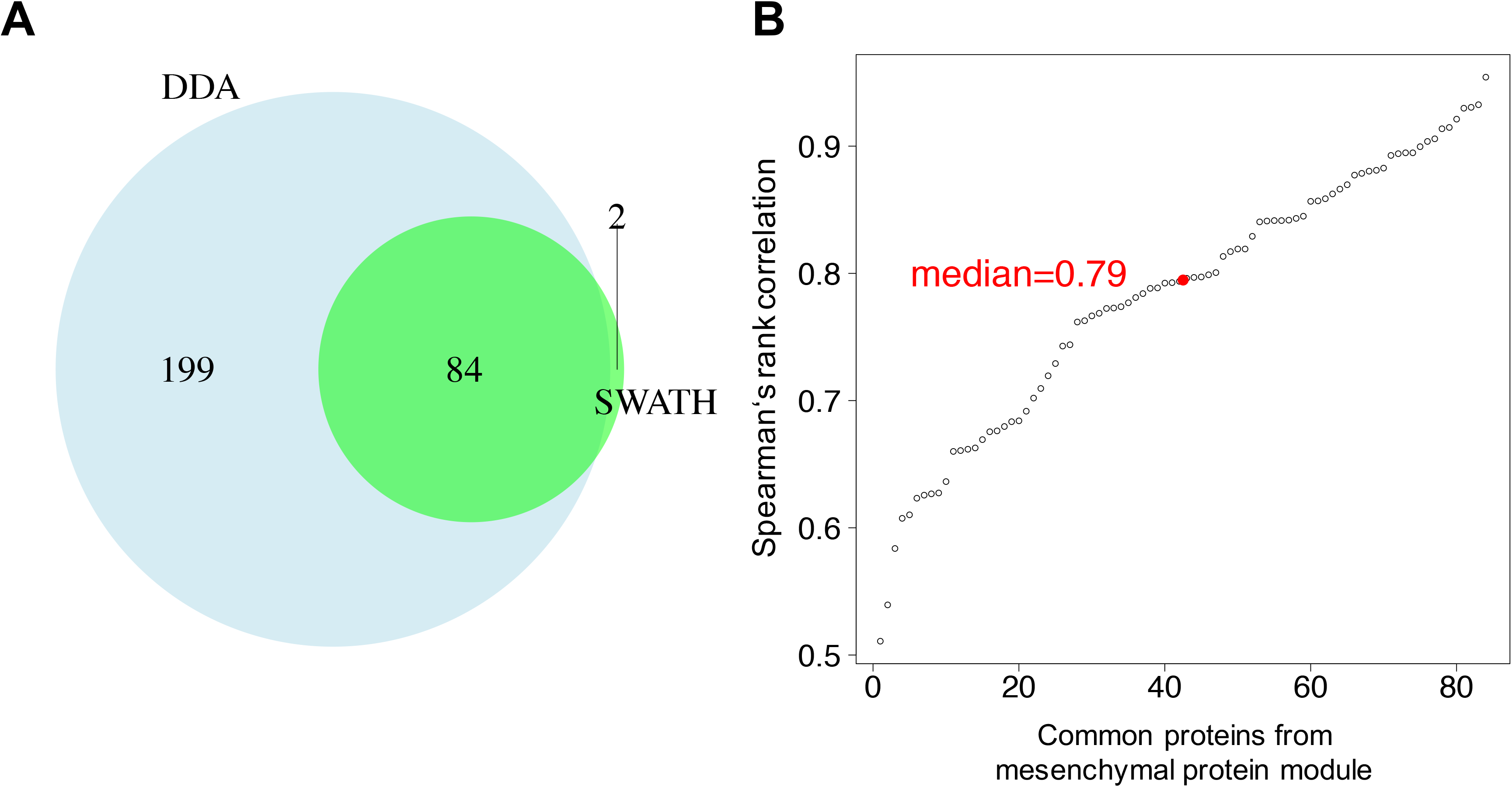
Comparison of proteins comprising the Mesenchymal subtype tumors in the iTRAQ DDA and SWATH-MS data. **a)** High overlap of proteins characterizing the Mesenchymal subtype in the SWATH MS data and the iTRAQ DDA data. **b)** Strong positive correlation of 84 proteins commonly comprising the Mesenchymal subtype in the iTRAQ DDA and SWATH-MS data (median *ρ* = 0.79).

To determine whether the same proteins from the iTRAQ DDA and SWATH-MS datasets would be identified in a group comparison of Mesenchymal subtype samples vs. the other samples, we used the sample subtype annotations from the Zhang *et al*. iTRAQ DDA study (Zhang et al., 2016a). The log_2_ fold-changes of these proteins in the iTRAQ DDA and SWATH-MS datasets were determined as well as the associated p-value (Figure 6). Proteins with significantly increased or decreased abundance (fold-change cut-off: 1.3; p<0.05) are indicated in red or blue, respectively. The significantly upregulated proteins (Figure 6A and **B, red dots**) in these comparisons showed a high overlap with the 84 common proteins from the ECM protein module (Figure 6C) as defined by the Zhang *et al*. iTRAQ DDA study (Zhang et al., 2016a). Well-established cancer markers such as Fibronectin 1 and Thrombospondins that are known for contributing to the metastatic progression of tumors (Hu et al., 2017; Incardona et al., 1993; Kenny et al., 2014; Mitra et al., 2011; Ricciardelli et al., 2016) were among those proteins, confirming the characterization of these samples as Mesenchymal subtype tumors. Although there is a high overlap of these proteins between the iTRAQ DDA and SWATH datasets, 33 of the up-regulated proteins were not identified as such in the iTRAQ DDA dataset. Nevertheless, most of those 33 proteins are part of the extracellular region and exhibit enrichment in extracellular matrix organization as well. Also of note, the quantitative dimension of iTRAQ DDA is narrower than that of the SWATH-MS data which influences the extent of similarity among the proteins quantified by each method.

**Figure 6.**
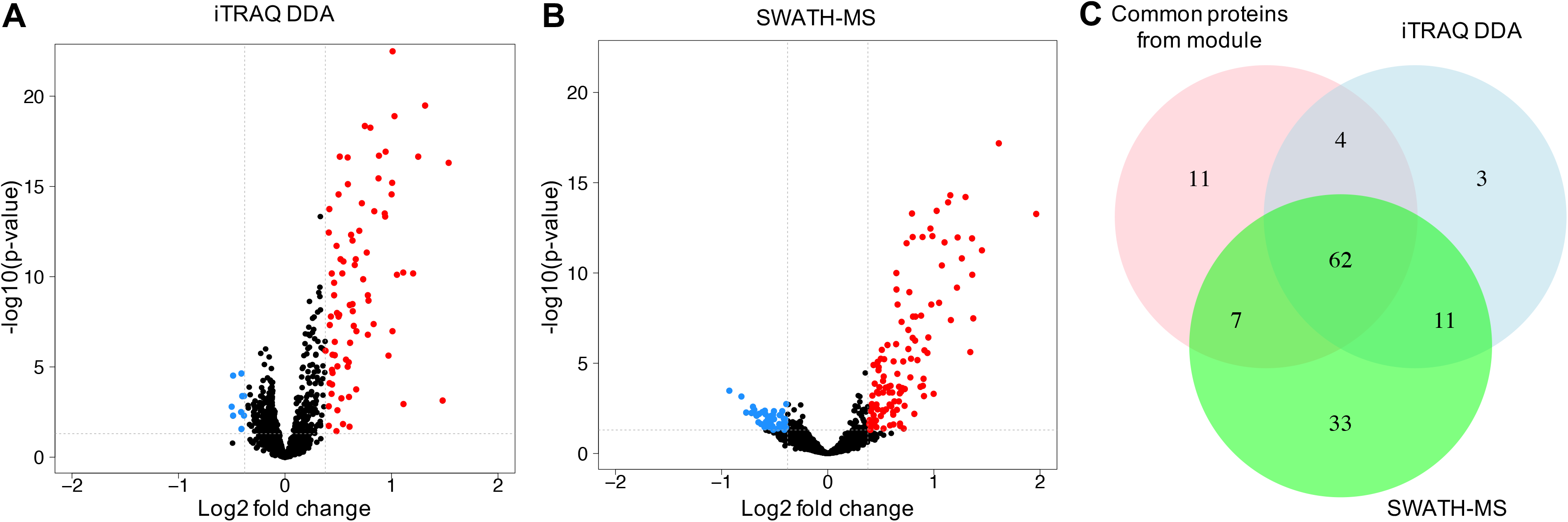
Group comparison of proteins in Mesenchymal subtype vs. the non-Mesenchymal subtypes. Up-regulated proteins among the Mesenchymal subtype tumors are indicated in red and the down-regulated proteins are indicated in blue. The fold-change cut-off was 1.3 and the significance cut-off was p<0.05. **a)** iTRAQ DDA data. **b)** SWATH-MS data. **c)** Comparison of all proteins assigned to the ECM module based on the original proteomic classification (Zhang et al., 2016a) vs. proteins assigned to the ECM module in the iTRAQ DDA data and the SWATH-MS data.

Since the proteomic subtypes other than the Mesenchymal subtype identified in the CPTAC and TCGA studies could not be clearly distinguished by the classification analyses of the data sets used in this study, we performed a similar differential expression analysis on those groups using the 1,599 proteins that were quantified in the iTRAQ DDA and SWATH-MS data sets (**Figure S5**). As expected, the p-values of those comparisons were considerably less significant than those resulting from the analyses of the Mesenchymal subtype. The Differentiated and Stromal subtypes did not lead to any conclusive results in these analyses based on the functional enrichment of the proteins with increased or decreased relative abundances, while the Proliferative and Immunoreactive subtypes were characterized by several differentially expressed proteins with rather high p-values (low significance). The WGCNA analyses of the respective differentially abundant proteins identified protein modules related to immune response and gene expression and translation (**Figure S4**). However, those modules did not show high associations with any of the patient groups.

### Homologous recombination deficiency (HRD)-related proteomic signature of HGSOC identified in SWATH-MS data

Previous studies have used gene expression and mutation profiles to characterize molecular subtypes of high-grade serous ovarian cancer to identify patients who respond well to poly-ADP ribose polymerase (PARP) inhibitor treatment (Lheureux et al., 2017; Tothill et al., 2008). Homologous recombination deficiency (HRD) is associated with a higher sensitivity towards PARP inhibitor treatment and therefore with a better prognosis for the respective patients. The initial iTRAQ DDA analysis of the HGSOC tumors resulted in the identification of a well-defined network of proteins with roles in histone acetylation that differentiated HRD from non-HRD tumors (Zhang et al., 2016a). Thus, we conducted an analysis to determine whether similar HRD-related features could be identified from the SWATH-MS data in the current study.

A group comparison of HRD vs. non-HRD patients using mapDIA (Teo et al., 2015) revealed several differentially expressed proteins (Figure 7A). Because none of these proteins could be directly linked to DNA repair mechanism-related functions, we used a network propagation approach (Hofree et al., 2013) to extend the comparatively limited analytical depth of the SWATH measurements. As an input we used a network obtained from STRING (Szklarczyk et al., 2015) filtered for highly confident experimental evidence (physical interactions with a score higher than 800) with 4,424 nodes. Log_2_-fold changes obtained from the group comparison approach were mapped on this network and these signals were propagated over the network to identify signals accumulating within a subnetwork. The top 5% (221 proteins) of the resulting positive and negative scores were used for further investigation and functional enrichment.

**Figure 7.**
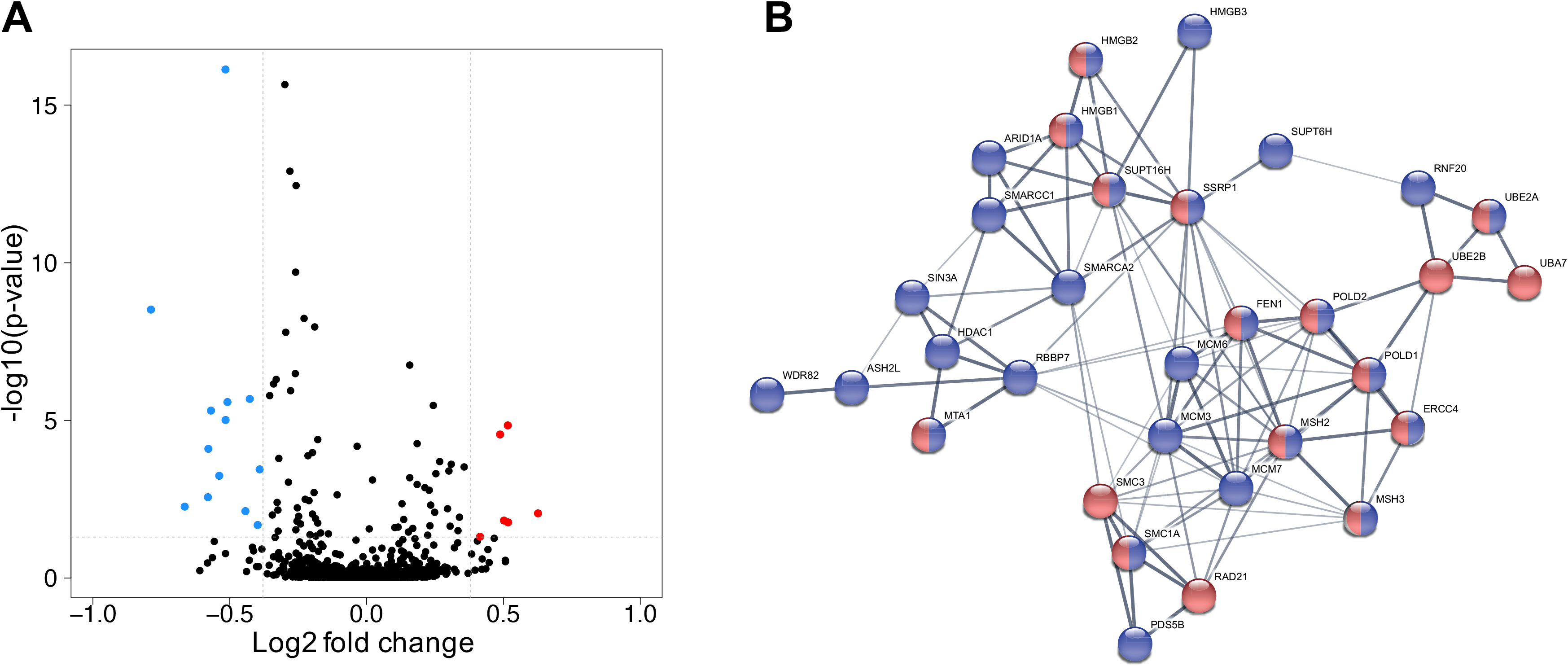
Group comparison of proteins in HRD vs. non-HRD tumors analyzed by SWATH-MS. **a)** Relative abundance of up- (red) and down-(blue) regulated proteins in the HRD compared to the non-HRD tumors. The log_2_ fold-change cut-off was 1.3 and the significance cut-off was p<0.05. **b)** Functional enrichment analysis using STRING. Blue nodes: Chromosome organization proteins. Red nodes: DNA repair proteins. Blue and red nodes: Proteins with roles in chromosome organization and DNA repair.

Further verifying that proteins with roles in histone acetylation differentiate HRD from non-HRD tumors (Zhang et al., 2016a), DNA repair and chromosome organization were among the GO terms that were identified using the previously described network propagation approach based on functional enrichment analysis. The respective sub-network contained proteins previously identified as belonging to the HRD-associated protein network, including histone deacetylase 1 (HDAC1; Figure 7B) and a histone-binding protein RBBP7 (Zhang et al., 2016a) (Figure 7B). HDAC1 and RBBP7 are among the blue nodes representing chromosome organization proteins. Another protein identified in this sub-network was a DNA mismatch repair protein which is a well-known marker for ovarian cancer, MSH2 (Maresca et al., 2015; Stewart et al., 2017; Xiao et al., 2014; Zhao et al., 2018). Additionally, subunits of the tumor suppressor complex SWI/SNF, including ARID1A, SMARCC1 and SMARCA2, were among the proteins in the enriched chromosome organization network. The interaction of SWI/SNF with PARP and BRCA1 has been demonstrated previously by *in vitro* studies using yeast two-hybrid screening, affinity purification followed by Western blotting and co-immunoprecipitation purification followed by LC-MS/MS (Bochar et al., 2000; Harte et al., 2010; Hill et al., 2004). The identification of SWI/SNF complex components in the enriched chromosome organization network in our study and the results from *in vitro* studies indicating the interaction of these proteins with PARP and BRCA1 support the well-known role of these proteins in the control of homologous recombination during DNA repair.

In summary, the analysis of the SWATH data was not only able to provide confirmation of some previously identified signatures from the iTRAQ DDA data in a larger untargeted network approach, but it also provided important mechanistic insights into HRD-related pathways as discovered by iTRAQ labeling and LC-MS/MS using DDA (Zhang et al., 2016a).

## Discussion

HGSOC is among the cancer types that have been proteogenomically characterized by CPTAC. The large-scale proteome analytical workflow employed by CPTAC uses isobaric tagging, offline peptide fractionation and LC-MS/MS. This workflow is considered a reference method for the comparison of tissue protein abundance across large sample cohorts. However, recently, DIA proteomic methods exemplified by SWATH-MS have been developed which are simpler, faster, cheaper, and consume less sample than the reference method, but provide shallower proteome coverage.

Although HGSOC is the most common histological subtype of ovarian cancer, there is a considerable amount of tumor heterogeneity (Arend et al., 2018), thus underscoring the need for comprehensive characterization of the molecular subtypes of this lethal disease. Previous studies have shown that somatic mutations (Wiegand et al., 2010), genetic (Gates et al., 2010) and environmental risk factors (Goode et al., 2010), and the clinical response rates to platinum- or taxane-based therapy (Sugiyama et al., 2000) vary considerably among the HGSOC molecular subtypes. As such, elucidating the molecular characteristics of HGSOC could facilitate the development of more targeted and effective therapies (Leong et al., 2015; Vaughan et al., 2011).

DDA mass spectrometry workflows have been employed to comprehensively characterize HGSOC (Coscia et al., 2016; Li et al., 2017a; Xie et al., 2017; Zhang et al., 2016a) with varying numbers of molecular subtypes emerging from these analyses. Functional genomic studies have identified various numbers of distinct HGSOC molecular subtypes with clinical relevance and pathways that are responsible for growth control in epithelial ovarian cancer (Cancer Genome Atlas Research, 2011; Tan et al., 2013). The discordance among these subtypes results from the varied sample sizes and analytical criteria used to conduct these studies (Helland et al., 2011; Tothill et al., 2008; Verhaak et al., 2013). Our current study provides orthogonal evidence of the proteomic signatures of the HGSOC Mesenchymal subtype and the fidelity of the HGSOC Mesenchymal subtype which is more sensitive to sample number compared to the number of quantified proteins. This has implications for the design of future large-scale clinical proteomic studies of cancer types where molecular subtyping is a predominant goal.

In this study, we compared the results obtained by the reference large-scale DDA proteomic method vs. SWATH-MS using aliquots of peptide samples generated for the CPTAC HGSOC study (Zhang et al., 2016a). The results indicate that iTRAQ DDA and SWATH-MS confidently identified differentially expressed proteins in enriched pathways associated with the Mesenchymal subtype of HGSOC tumors as evidenced by the: 1) high degree of overlap of up-regulated extracellular matrix-related proteins (Figure 5A), 2) strongly positive median correlation (*ρ* = 0.79) among the proteins comprising this subtype from both proteomic workflows (Figure 5B), and 3) statistically significant stability of this subtype in the context of the number of tumors included in the clustering analysis (**Figure S3B**). The robustness of the Mesenchymal subtype with respect to molecular subtype cluster stability could be a signature of the decreased survival of patients whose tumor samples express this molecular signature compared to an Immunoreactive signature. A study conducted using a cohort of 174 HGSOC patients from the Mayo Clinic with long-term clinical follow-up observed statistically significantly worse survival of patients whose tumor samples expressed a Mesenchymal-like signature upon the analysis of a set of 1,850 genes (Konecny et al., 2014). A similar trend of worse survival for patients with Mesenchymal subtype tumors compared to those with Immunoreactive subtype tumors was observed upon the analysis of a separate cohort of 185 HGSOC patients (Bonome et al., 2008).

The notion of discrete HGSOC subtypes that are mutually exclusive is not universally accepted. Verhaak *et al*. suggested that an individual tumor could be represented by multiple signatures based on different levels of pathway activation (Verhaak et al., 2013). This concept has been supported by Konecny *et al*. who proposed a multi-dimensional approach to subtyping where molecular subtypes lie on a spectrum with partly overlapping causes (Konecny et al., 2014). Additional large-scale clinical proteomic studies of HGSOC tumors that are designed to address issues related to tumor heterogeneity would be beneficial in enhancing the resolution of molecular subtyping.

Our SWATH-MS analysis confirmed the proteomic signature of HRD established by iTRAQ DDA analysis wherein a sub-network of *BRCA1*- or *BRCA2*-related proteins displayed co-expression patterns differentiating HRD from non-HRD patients (Zhang et al., 2016a). Many of the proteins in these identified modules have roles in histone acetylation or deacetylation. Inhibitors of PARP and histone deacetylase inhibitors have emerged as novel classes of anti-cancer drugs to treat HR-related ovarian cancer associated with *BRCA1/2* mutations (Bryant et al., 2005; Farmer et al., 2005; Yano et al., 2018; Yuan et al., 2017). Thus, the proteomic characterization of HGSOC tumor tissue biopsies from patients who receive PARP inhibitor treatment could be beneficial in further elucidating the molecular mechanisms that are implicated in HRD.

One of the strengths of the iTRAQ DDA workflow is the ability to achieve deep proteome coverage which exceeds that of the SWATH workflow by almost 3-fold. A total of 8,597 proteins were quantified by iTRAQ DDA compared to 2,914 proteins by SWATH-MS. These numbers represent the aggregate numbers of quantified proteins. However, after employing filtering strategies to restrict the data to only the proteins that were quantified across all 103 tumors for each workflow, these numbers decreased to 4,363 and 1,659, respectively. The group of 1,599 proteins that were quantified by both proteomic workflows was used to compare the performance of iTRAQ DDA and SWATH-MS.

In addition to their analytical performance, there are also considerable differences in the resource characteristics of iTRAQ DDA and SWATH-MS, including sample requirement, sample throughput and cost. Hence, it is clear that iTRAQ DDA for comprehensive proteome profiling is a substantially more resource-intense workflow compared to SWATH-MS.

Based on the concordance between the iTRAQ DDA and SWATH-MS results that we have shown in this study, SWATH-MS, which is considerably less resource-intense than iTRAQ DDA, can be reliably deployed in the proteomic analysis of clinical specimens. The clinical utility of future large-scale translational proteomic studies, regardless of the employed analytical methodology, can be strengthened by the use of large sample cohorts that have undergone comprehensive pathology review.

Although our study was focused on the performance of SWATH-MS, other DIA methods are currently available. Our SWATH-MS analysis was conducted using a 5600^+^ Triple-TOF mass spectrometer; however, a newer generation of this mass spectrometer platform exists with an increased linear dynamic range and enhanced detection system, which could result in improved instrument performance with respect to the number of quantifiable proteins per analytical run. A recent DIA study conducted by Bruderer *et al*. using a quadrupole ultra-high field Orbitrap mass spectrometer resulted in the identification of more than 6,000 proteins in human cell lines and more than 7,000 proteins in mouse tissues (Bruderer et al., 2017). Of note, the use of isobaric chemical labeling strategies, including 10- and 11-plex tandem mass tags (TMT), has greatly facilitated the multiplexing capabilities of large-scale clinical proteomic studies resulting in the reproducible quantification of >10,000 proteins (Krieger et al., 2019; Mertins et al., 2018).

In contrast to the original CPTAC HGSOC study, we did not assess protein phosphorylation in the current study. However, it should be noted that SWATH/DIA-MS has been shown to be compatible with the analysis of protein phosphorylation patterns, and specific software tools supporting such analyses have been developed (Rosenberger et al., 2017b).

As expected, the development of new analytical instruments and methods often enables an expanded breadth and/or depth of analytical measurement. After the performance of these instruments and methods has been optimized and validated, the short-comings of previously existing methods become evident. It is as-yet unknown whether SWATH-MS and other DIA proteomic methods will have an increased prevalence in clinical proteomic analyses. However, our current study provides compelling orthogonal evidence that SWATH-MS elucidates some of the common biological signatures of the Mesenchymal subtype of HGSOC.

## Acknowledgments

We acknowledge contributions from Vlad Petyuk who provided scripts and methods from the previously published CPTAC study. We also thank Ludovic Gillet, Isabell Bludau, Ben Collins and the computational proteomics team at ETH Zürich for their important contributions in discussions. We are grateful to Sciex (Christie Hunter, Mark Cafazzo and George Manning) for the 5600^+^ TripleTOF mass spectrometer and access to open-source SWATH data processing software. Funding for the study was provided by the National Cancer Institute Clinical Proteomic Tumor Analysis Consortium (NCI CPTAC, U24CA160036 and U24CA210985 to H.Z. and D.W.C.), the European Research Council (AdG-233226 Proteomics v.3.0 and AdG-670821 Proteomics 4D to R.A.) and the Swiss National Science Foundation (grant 31003A_166435 to R.A.).

## Author Contributions

SNT performed the experiments with sample preparation and SWATH-MS analysis; BF conducted computational analyses for SWATH data and bioinformatics analysis; SNT and MS performed and assisted in computational analyses with iTRAQ DDA data. RA, HZ, and DWC conceived the study, designed the experiments, and supervised the participants. All authors participated in the preparation of the manuscript.

## STAR Methods

### KEY RESOURCES TABLE

| REAGENT or RESOURCE | SOURCE | IDENTIFIER |
| --- | --- | --- |
| **Biological Samples** |  |  |
| Human ovarian tumor tissue | National Cancer Institute Clinical Proteomic Tumor Analysis Consortium | https://proteomics.cancer.gov/programs/cptac |
| **Chemicals, Peptides, and Recombinant Proteins** |  |  |
| Urea | Sigma-Aldrich | Cat# U0631 |
| HPLC-grade water | J.T. Baker | Cat# 4218-03 |
| Triethylammonium bicarbonate | Sigma-Aldrich | Cat# T7408 |
| Ammonium bicarbonate | Sigma-Aldrich | Cat# A6141 |
| Tris (2-carboxyethyl) phosphine (TCEP) | Thermo Fisher Scientific | Cat# 20490 |
| Iodoacetamide | Sigma-Aldrich | Cat# A3221 |
| Sequencing-grade modified trypsin | Promega | Cat# V511X |
| Formic acid | Sigma-Aldrich | Cat# 56302 |
| Trifluoroacetic acid | Sigma-Aldrich | Cat# 91707 |
| Acetonitrile | J.T. Baker | Cat# 9829-03 |
| iTRAQ reagent kit – 4plex | Sciex | Cat# 4352135 |
| **Critical Commercial Assays** |  |  |
| BCA bicinchoninic protein assay kit | Pierce | Cat# 23325 |
| **Deposited Data** |  |  |
| iTRAQ DDA data | NCI CPTAC Data Portal | https://cptac-data-portal.georgetown.edu/cptac/s/S026<br>(Mass spectrometry site: Johns Hopkins University) |
| SWATH data, input spectral library, OpenSWATH results | ProteomeXchange Consortium | http://proteomecentral.proteomexchange.org<br>Dataset identifier: PXD010437 |
| **Software and Algorithms** |  |  |
| Programming code for bioinformatics pipeline | Github repository | [bfriedrichgrube/OC_CPTAC_iTRAQ_SWATH](https://github.com/bfriedrichgrube/OC_CPTAC_iTRAQ_SWATH) |
| msconvert | ProteoWizard v3.0.8851 | http://proteowizard.sourceforge.net/download.html |
| X!Tandem 2013.06.25.1 | Duncan et al., 2005 | https://www.thegpm.org/tandem/ |
| OMSSA 2.1.9 | Geer et al., 2004 | https://omictools.com/omssa-tool |
| Comet 2015.02 rev. 3 | Eng et al., 2013 | http://comet-ms.sourceforge.net/ |
| Trans-Proteomic Pipeline v. 4.7 | Keller et al., 2005;<br>Lam et al., 2007 | http://tools.proteomecenter.org/wiki/index.php?title=Software:TPP |
| OpenSWATH | Rost et al., 2014 | http://www.openswath.org/en/latest/ |
| SpectraST | Keller et al., 2005;<br>Lam et al., 2007 | http://tools.proteomecenter.org/wiki/index.php?title=SpectraST |
| TRIC | Rost et al., 2016 | http://proteomics.ethz.ch/tric/ |
| mapDIA v. 3.0.2 | Teo et al., 2015 | http://mapdia.sourceforge.net/Main.html |
| **Other** |  |  |
| SepPak tC18 Vac cartridges | Waters | Cat# WAT054925 |
| Polysulfoethyl A strong cation exchange columns | Glygen | Cat# TT3SSA |

### CONTACT FOR REAGENT AND RESOURCE SHARING

Further information and requests for resources and reagents should be directed to and will be fulfilled by the Lead Contact, Hui Zhang.

### EXPERIMENTAL MODEL AND SUBJECT DETAILS

### Tumor samples

The tumor specimens were obtained through The Cancer Genome Atlas (TCGA) Biospecimen Core Resource, and they were previously genomically characterized (Cancer Genome Atlas Research, 2011). As previously described (Zhang et al., 2016a), the biospecimens were obtained from newly diagnosed patients with ovarian serous adenocarcinoma who were undergoing surgical resection and did not receive prior treatment, including chemotherapy or radiotherapy, for their disease. Frozen tissue specimens were extracted and used for subsequent proteomic analysis.

## METHOD DETAILS

### Protein extraction and in-solution digestion

Approximately 50mg of each tumor tissue specimen was sonicated in 1.5mL of 8M urea, 0.8M NH_4_HCO_3_, pH 8.0. Protein concentration was determined using a BCA assay (Thermo Fisher Scientific). Protein disulfide bonds were reduced with 10mM tris (2-carboxyethyl) phosphine (TCEP) for 1h at 37°C, followed by alkylation with 12mM iodoacetamide for 1h at RT in the dark. After dilution 1:4 with deionized water, proteins were digested with sequencing-grade modified trypsin (Promega, Madison, WI) (1:50 enzyme:protein, weight/weight) for 12h at 37°C. This was followed by the addition of an aliquot of the same amount of trypsin and incubation overnight at 37°C. The digested samples were acidified with 10% trifluoroacetic acid (TFA) to pH<3, de-salted using strong cation exchange and C18 solid-phase extraction (SPE) columns (Waters, Milford, MA) and dried using a Speed-Vac.

### Shotgun proteomics using iTRAQ data-dependent acquisition (DDA)

Relative quantitative proteomic analysis was conducted using 4-plex isobaric tags for relative and absolute quantitation (iTRAQ) reagents (Sciex) as previously described (Zhang et al., 2016a). Peptides (500µg) were dissolved in 150µL of 0.5M triethylammonium bicarbonate, pH 8.5 and combined with 5U of 4-plex iTRAQ reagent dissolved in ethanol followed by a 2h incubation at RT, quenching with 10% TFA, and de-salting using C18 SPE columns. iTRAQ channel 114 was used to label the reference sample which was created by pooling an aliquot from each individual tumor sample. Offline basic reversed phase liquid chromatography (bRPLC) was conducted using a Zorbax extend 4.6 x 100mm C-18 column (Agilent) and an Agilent 1220 Infinity HPLC system to reduce the sample complexity prior to mass spectrometry analysis. A total of 96 fractions were collected and concatenated into 24 fractions. The fractions were dried in a Speed-Vac and stored at −80°C until analysis by LC-MS/MS using an LTQ Orbitrap Velos mass spectrometer (Thermo Scientific). Peptides were loaded onto a 2cm guard column (Thermo Scientific) and separated on a 75µm x 15cm Acclaim PepMap100 column (Thermo Scientific) using a Dionex Ultimate 3000 RSLC nano chromatography system (Thermo Scientific). The LC gradient profile was 2-22% B for 70min, 22-29% B for 8min, 29-95% B for 4min, and 95% B for 8min, where mobile phase B was 0.1% formic acid in acetonitrile, and mobile phase A was 0.1% formic acid in water. Orbitrap full MS spectra were collected from 400-1800 m/z at a resolution of 30,000 followed by data-dependent MS/MS (7,500 resolution) of the ten most abundant ions. Charge-state screening was enabled to prevent the acquisition of MS/MS spectra for ions with unassigned or single charges. Dynamic exclusion (40s duration) was enabled to minimize the repeated acquisition of previously acquired MS/MS spectra.

### Proteomic analysis using SWATH mass spectrometry

SWATH-MS measurements were conducted at the Johns Hopkins University using a Sciex 5600^+^ TripleTOF mass spectrometer interfaced with an Eksigent ekspert nanoLC 425 cHiPLC system. Peptides (1µg) were loaded onto a 6mm x 200µm ChromXP C18-CL 3µm, 120Å trap column followed by separation on a 75µm x 15cm ChromXP C18-CL 3µm, 120Å Nano cHiPLC column using a 120min method (90min gradient from 3-35% B – 0.1% formic acid in acetonitrile) at a flow rate of 300 nL/min. To create the spectral library for the SWATH-MS data analysis, each sample was run individually (1µg peptides per injection) using a data-dependent data acquisition (DDA) method wherein MS spectra were acquired across a range of 400-1800 m/z followed by the acquisition of MS/MS spectra of the top 30 most intense precursor ions with a charge state of z=2-5. The spectral library was also comprised of mass spectrometry data acquired from a fractionated (48 fractions) pool of peptides from all 103 tumors. Each of the 48 fractions from the pooled sample was analyzed using the same DDA method described above. SWATH data of the individual tumors were acquired using a variable window strategy wherein the sizes of the precursor ion selection windows were inversely related to m/z density. The average window width for precursor ion selection was 12 m/z with a range of 6-25 m/z. The collision energy was optimized for each window according to the calculation for a charge 2+ ion centered in the window with a spread of 5 eV. The MS accumulation time was 250ms and the MS/MS accumulation time for fragment ions accumulated in high sensitivity mode was 50ms, resulting in a total duty cycle of approximately 3.5s. To assess the analytical precision of the proteomics measurements, peptides from trypsin-digested HEK293 cell proteins were analyzed via SWATH-MS in triplicate immediately prior to the SWATH-MS analysis of the 103 individual tumor samples. Additionally, an ovarian cancer tumor sample separate from the TCGA collection was used as a QC sample. This specimen was analyzed using the DDA method described above in duplicate prior to the SWATH-MS analysis of the 103 individual tumor samples and 10 days later in duplicate mid-way through the analysis of the individual tumor samples. Instrument performance was assessed daily by monitoring the peak area of 5 peptides from a trypsin-digested *E. coli* ß-Galactosidase LC-MS standard that was injected every day a sample was run on the mass spectrometer.

### iTRAQ DDA data processing

iTRAQ DDA data from the Johns Hopkins CPTAC center was analyzed by the Common Data Analysis Pipeline (CDAP) and downloaded from the NCI CPTAC Data Portal (https://cptac-data-portal.georgetown.edu/cptac/s/S026); mass spectrometry site: Johns Hopkins University (Edwards et al., 2015; Ellis et al., 2013). Values obtained from proteotypic peptides were chosen for further analysis. The protein matrix contained 8597 proteins. The protein matrix was filtered for proteins without missing values (4363 proteins). PSM files were additionally downloaded from the CPTAC Data Portal and processed following the CDAP to the peptide level data in order to investigate peptide variability.

### SWATH-MS data processing

Raw mass spectrometry measurements obtained from the TripleTOF 5600^+^ in DDA and SWATH mode were converted to mzXML file format using msconvert (ProteoWizard v3.0.8851) (Chambers et al., 2012). DDA measurements from all 103 samples and 48 fractions of a pooled sample were searched with X!Tandem (2013.06.25.1) (Duncan et al., 2005), OMSSA (2.1.9) (Geer et al., 2004) and Comet (2015.02 rev. 3) (Eng et al., 2013). Identified peptides were processed through the Trans-Proteomic Pipeline (TPP v.4.7 Polar Vortex rev 0) using PeptideProphet, iProphet and ProteinProphet scoring (FDR <0.01) and SpectraST (Keller et al., 2005; Lam et al., 2007). The assay library was built according to (Schubert et al., 2015) using a set of 113 common internal retention time standards (ciRTs) (Parker et al., 2015). The resulting library contained 8897 proteins supported by proteotypic peptides.

SWATH data was analyzed using OpenSWATH (Rost et al., 2014) from OpenMS (v.1.10.0) with the previously described sample-specific assay library. FDR was controlled using PyProphet (v.0.0.19) allowing for a peptide-query FDR of 1% and protein-FDR of 1% as described by Rosenberger et al. (Rosenberger et al., 2017a). Runs were aligned for improved quantification using TRIC (msproteomicstools master branch from GitHub commit c10a2b8) (Rost et al., 2016).

OpenSWATH results were annotated and formatted using the R package SWATH2stats (v.1.0.3) (Blattmann et al., 2016) prior to quantile normalization (preprocessCore v.1.32.0) (Bolstad, 2017) and batch-wise mean-centering batch correction of known sample preparation batches. Fragment-level data was then fed into mapDIA (v.3.0.2) (Teo et al., 2015) in order to filter for outliers and proteins with at least 3 fragments per peptide and 2 peptides per protein. The 3 most intense fragments per peptide were summed and the resulting peptide intensities were divided by the mean peptide intensity (as an internal reference) over all runs to obtain a similar data structure as in iTRAQ DDA. Peptide ratios were averaged per protein. A total of 2914 proteins were quantified in this manner across all 103 samples.

An additional filtering and imputation step was conducted to mitigate the missing values in the SWATH data. **Figure S2B** shows the number of missing values in the SWATH data and their occurrence dependent on the protein abundance represented by log_2_ intensity (**Figure S2C**). As expected, there is an inverse relationship between protein abundance and missing values. Missing values in low abundance proteins are more likely to occur due to technical reasons, while missing values amongst high abundance proteins potentially have biological significance which might be important for further analysis. We therefore filtered the protein matrix based on the occurrence of abundance-dependent missing values. We confined the number of missing values using a stricter threshold for low abundance proteins than high abundance proteins according to the following rules: no missing values were allowed for the lowest abundant 10% of proteins, 10% missing values were allowed for the next 10% abundant proteins and so on until 90% missing values were allowed for the highest abundant proteins. A total of 1659 proteins remained passed these filtering steps. The missing values from those proteins were filled using an imputation method based on what is used by Perseus software (Tyanova et al., 2016) wherein random numbers forming a normal distribution were generated fitting in the left end of the distribution of the measured values, representing values within the noise.

## QUANTIFICATION AND STATISTICAL ANALYSIS

### Bioinformatics pipeline

Bioinformatic analyses were performed in R (v.3.2.2 “Fire Safety”).

#### Technical evaluation

A total of 1599 proteins common to the iTRAQ DDA and SWATH data sets were evaluated for their correlation and variability (Figure 2). The distribution of values in both data sets was plotted using the R package vioplot (v.0.2) (Adler, 2005). Spearman’s rank correlation was computed for each protein across all 103 samples in both data sets. The resulting Spearman’s rho values were ordered and the median was calculated. For peptide variability, a CV-like score was calculated for the peptides from each protein (standard deviation of peptides per protein divided by mean peptide intensity per protein). The distribution of the resulting CVs was plotted with vioplot.

#### Molecular classification comparison

Protein matrices were transformed into z-scores and a molecular classification was obtained by running a model-based clustering with mclust from the R package mclust under default settings (v.5.2) (Fraley and Raftery, 2002; Fraley et al., 2012). Protein modules characterizing sample groups were obtained with weighted gene-correlation network analysis (WGCNA v.1.51) (Langfelder and Horvath, 2008, 2012) and functional enrichment with clusterProfiler (v.2.4.3) (Yu et al., 2012) (**Figure S4**). The resulting molecular classifications were compared based on their similarity using the Adjusted Rand Index (ARI), a measure for classification similarity, implemented within the R package mclust previously used. The heat map displayed in **Figure S3A** was generated based on SWATH z-scores using pheatmap (v.1.0.8) (Kolde, 2015). The stability of the classification results was evaluated by bootstrapping. Different fractions of samples or proteins were drawn 100 times and classified using mclust with default settings, and the similarity to the molecular classification of the full data set was compared with the ARI (Figure 4).

In order to assess the stability of the mesenchymal class a Fisher’s exact test was performed on the mesenchymal class and the other resulting classes with a bootstrapping approach by drawing a fraction of samples 100 times (**Figure S3B**). Group comparison of the original classes from CPTAC (Zhang et al., 2016a) were obtained by a student’s t-test (Figure 6 and **S4**).

#### HRD group comparison

Group comparison to identify the differentially expressed proteins in the HRD vs. non-HRD patients was conducted using mapDIA (v.3.0.2) (Teo et al., 2015) on fragment ion-level data (Figure 7A). The resulting fold changes were used as input signal in a network propagation approach (Hofree et al., 2013) called Network Smoothing (R package BioNetSmooth v.1.0.0) (Chokkalingam et al., 2016).

As an input network the STRING database for human (Taxon 9606 v.10.5) was downloaded and filtered for interactions with experimental evidence and a score >800. The top 5% of negative and positive scores were used for further investigation of functional enrichment of Gene Ontology using the STRING database (STRING-db.org v.10.5 accessed on 2018/03/14).

## DATA AND SOFTWARE AVAILABILITY

### Mass spectrometry data

All the raw data from SWATH-MS measurements, along with the input spectral library and OpenSWATH results can be freely downloaded from the ProteomeXchange Consortium (http://proteomecentral.proteomexchange.org) with the dataset identifier: PXD010437 via the PRIDE partner (Vizcaino et al., 2016) (Reviewer account details: Username, Password: 9VoPWx2g). The iTRAQ DDA data are available on the NCI CPTAC Data Portal (https://cptac-data-portal.georgetown.edu/cptac/s/S026); mass spectrometry site: Johns Hopkins University. Other data are available from the corresponding authors upon request.

### Programming codes

All code used for the downstream bioinformatics pipeline can be obtained from the github repository (bfriedrichgrube/OC_CPTAC_iTRAQ_SWATH).

## Supplemental Information

**Figure S1. Distribution of quantified proteins from QC strategies used to assess technical variability. a)** Distribution of quantified protein CVs from HEK293 lysate analyzed via SWATH (n=3 technical replicates per day for 3 days; total = 9 technical replicates) prior to the analysis of the ovarian tumors. Median total CV=15%; 3,855 quantified proteins. **b)** Distribution of quantified protein CVs from a control ovarian tumor analyzed in duplicate via DDA prior to and mid-way through the ovarian tumor SWATH analyses (n=4 technical replicates). Median total CV=7%; 781 quantified proteins.

**Figure S2. Completeness of iTRAQ DDA and SWATH-MS proteomic data. a)** Analysis of missing values in the iTRAQ DDA dataset. **b)** Analysis of missing values in the SWATH-MS dataset. **c)** Relationship between signal intensity and number of missing values for the proteins quantified by SWATH-MS.

**Figure S3. Protein (iTRAQ DDA, SWATH-MS) and mRNA-based high-grade serous ovarian cancer subtype classification. a)** Tumor subtype classification based on the associated driving protein modules. Numbers indicate z-scores. Subtype classification: Blue – Mesenchymal; Red – Differentiated; Purple – Proliferative; Green – Immunoreactive; Yellow – Stromal. **b)** Stability of the Mesenchymal compared to the non-Mesenchymal subtypes assessed by bootstrapping and Fisher’s exact test.

**Figure S4. Analysis of tumor subtype analysis according to data type.** Correlation of weighted gene co-expression network analysis (WGCNA)-derived protein modules with proteomic subtypes derived from the iTRAQ DDA data (**a**) and the SWATH-MS data (**b**). The numbers in each box indicate the Pearson correlation coefficient (−1 to 1), and the numbers in parentheses indicate the p-value corresponding to the significance of each association. The enrichment of Reactome pathways in the WGCNA-derived modules in the iTRAQ DDA and SWATH-MS data is also shown. The significance of association is denoted by colored dots indicating the q-value, and the number of proteins mapped to a given ontology as a proportion of the total number of proteins in the module (GeneRatio) is indicated by the size of the dot. The WGCNA-derived protein modules for the iTRAQ DDA data are as follows: magenta – gene expression; black – immune response; purple – transcription; tan – erythrocyte and platelet; magenta – cytokine signaling; red – ECM interaction; gray – metabolism. The WGCNA-derived protein modules for the SWATH data are as follows: green – immune response; turquoise – gene expression; black – ECM interaction; red – complement cascade; gray – metabolism.

**Figure S5. Differential relative protein abundance analysis of proteins in the Differentiated, Immunoreactive, Proliferative and Stromal subtype tumors from the iTRAQ DDA and SWATH-MS datasets.** Up-regulated proteins are indicated in red and the down-regulated proteins are indicated in blue. The log_2_ fold-change cut-off was 1.3 and the significance cut-off was set at p<0.05.

