## Supplementary Figures for "Orthogonal proteomic platforms and their implications for the stable classification of high-grade serous ovarian cancer subtypes"

Supplementary Figure 1: Distribution of quantified proteins from QC strategies used to assess technical variability

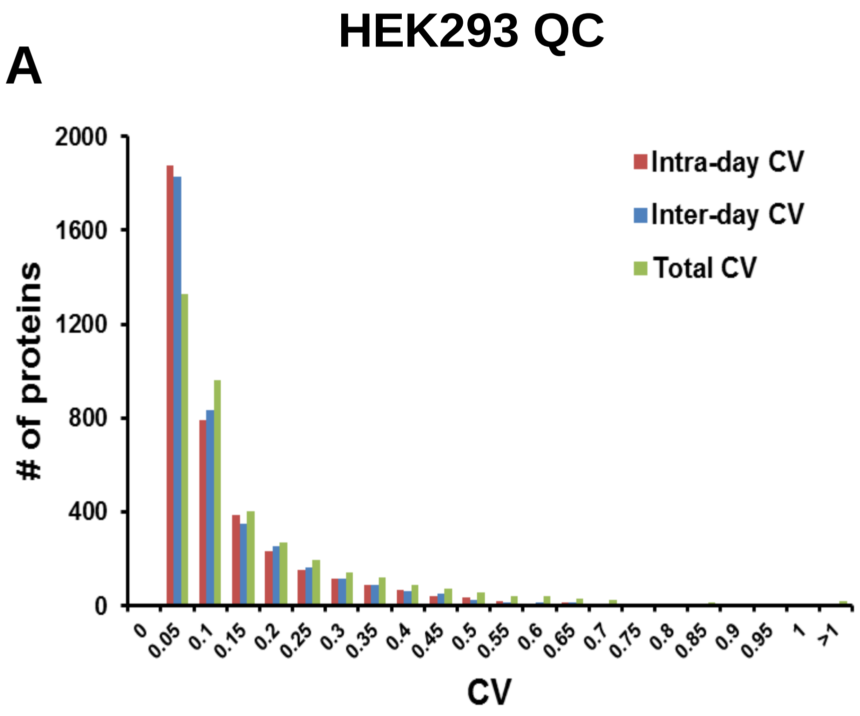


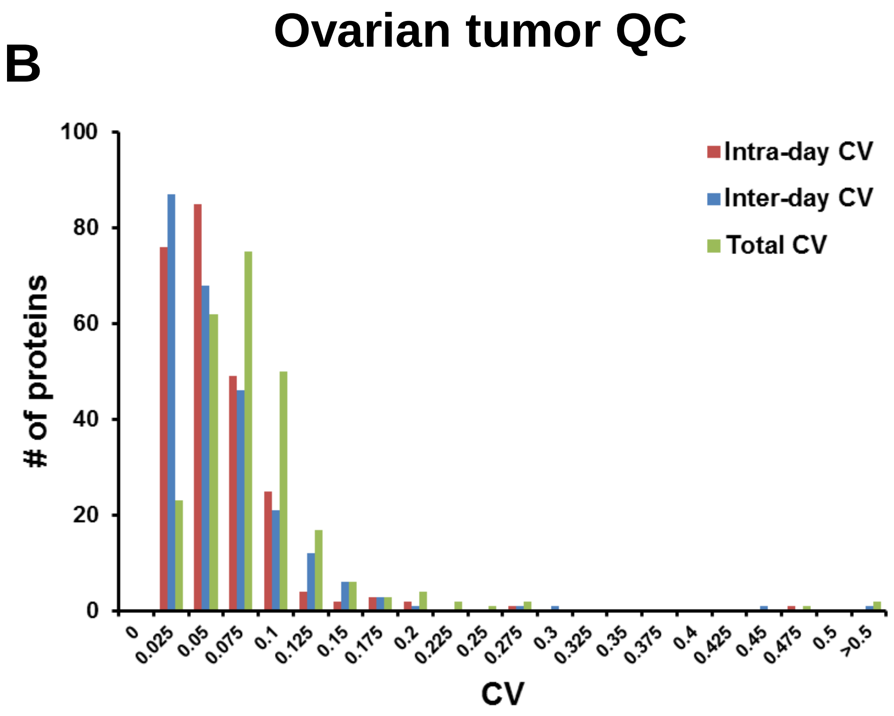

### Supplementary Figure 2: Completeness of iTRAQ DDA and SWATH-MS proteomic data

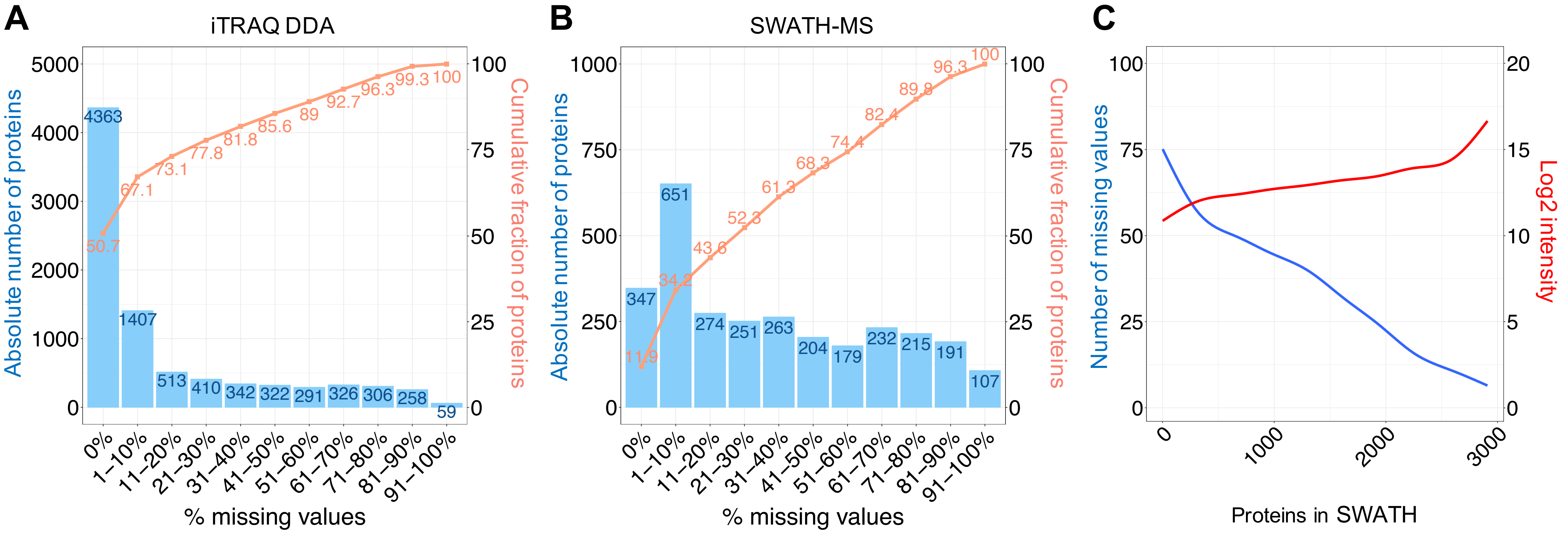

Supplementary Figure 3: Protein (iTRAQ DDA, SWATH-MS) and mRNA-based high-grade serous ovarian cancer subtype classification

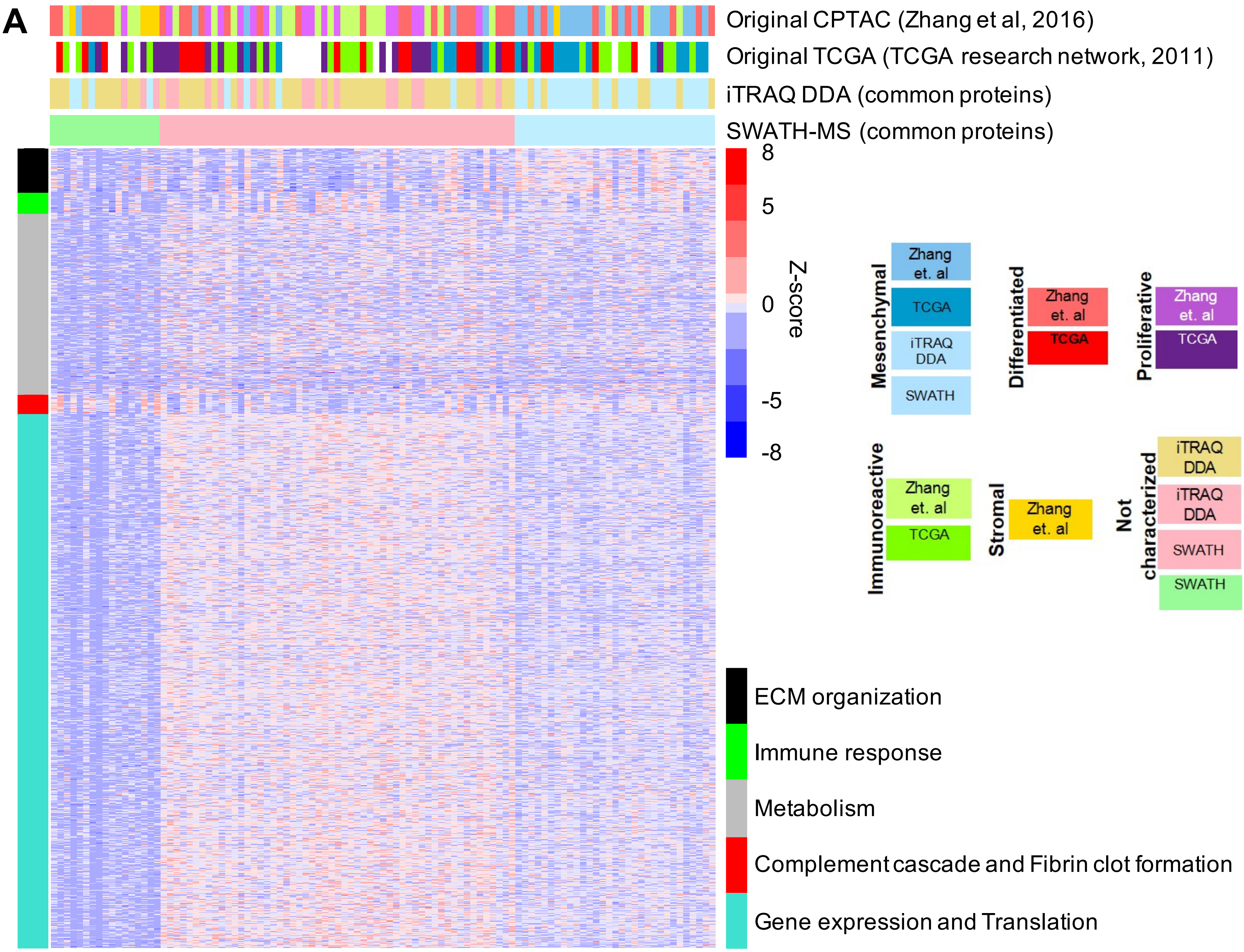


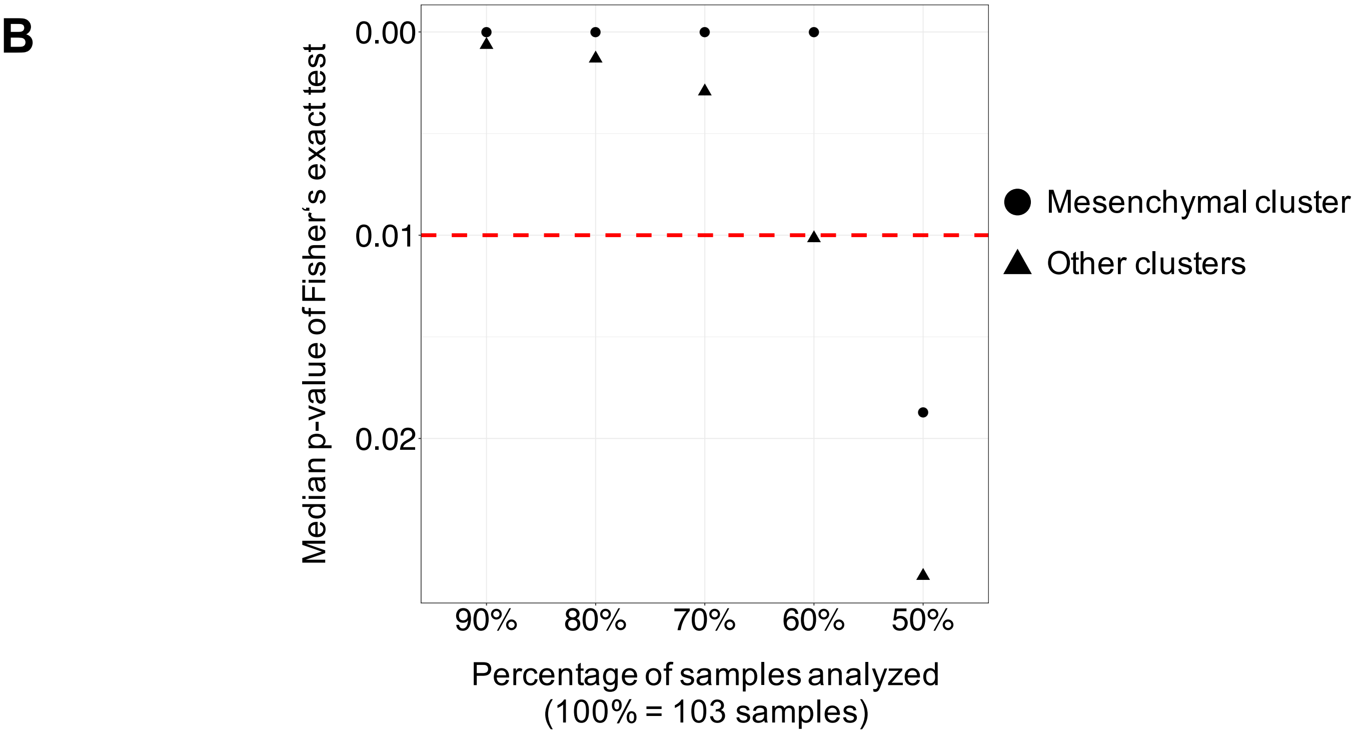

Supplementary Figure 4: Analysis of tumor subtype according to data type

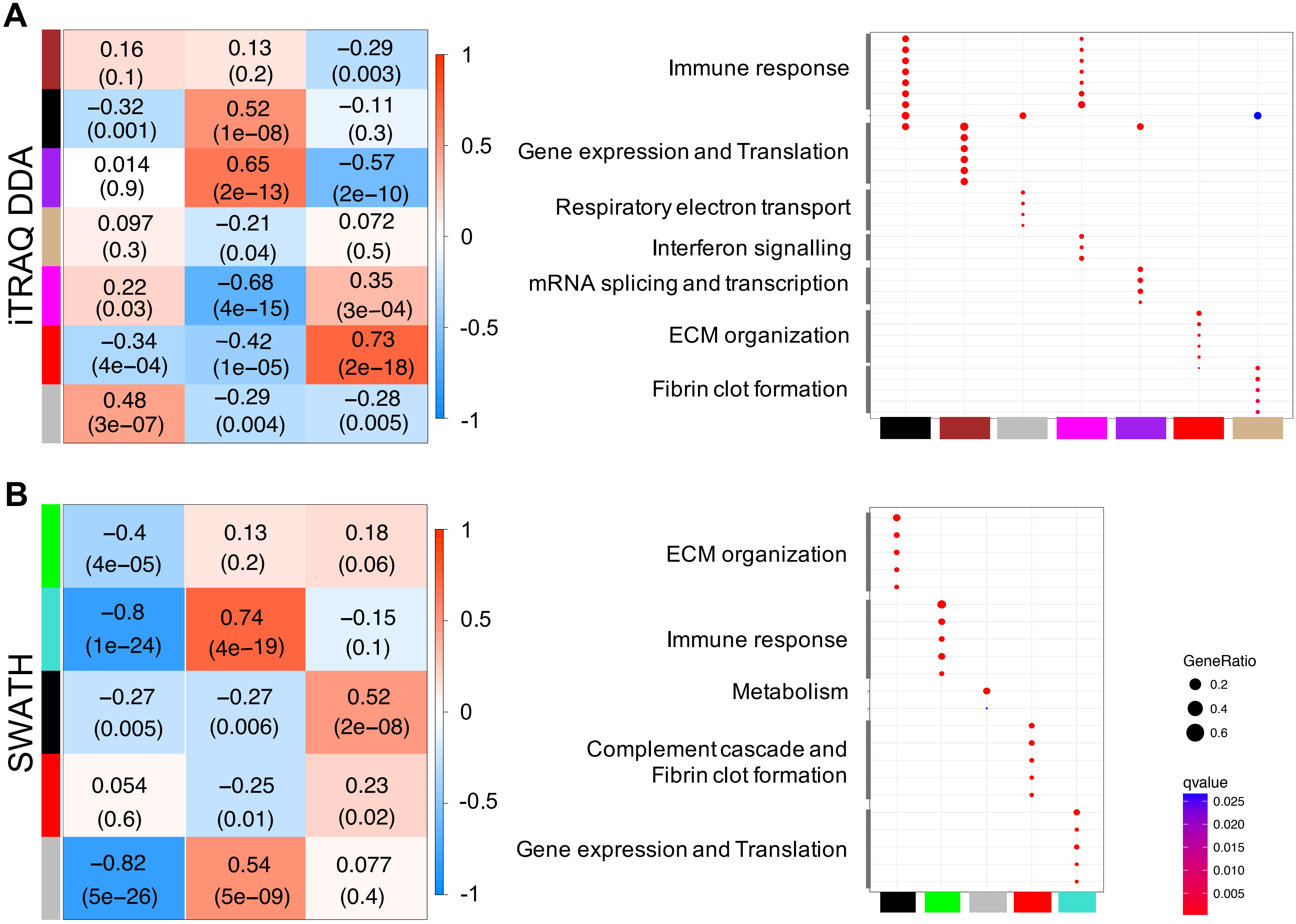

Supplementary Figure 5: Differential relative protein abundance analysis of proteins in the Differentiated, Immunoreactive, Proliferative and Stromal subtype tumors from the iTRAQ DDA and SWATH-MS datasets.

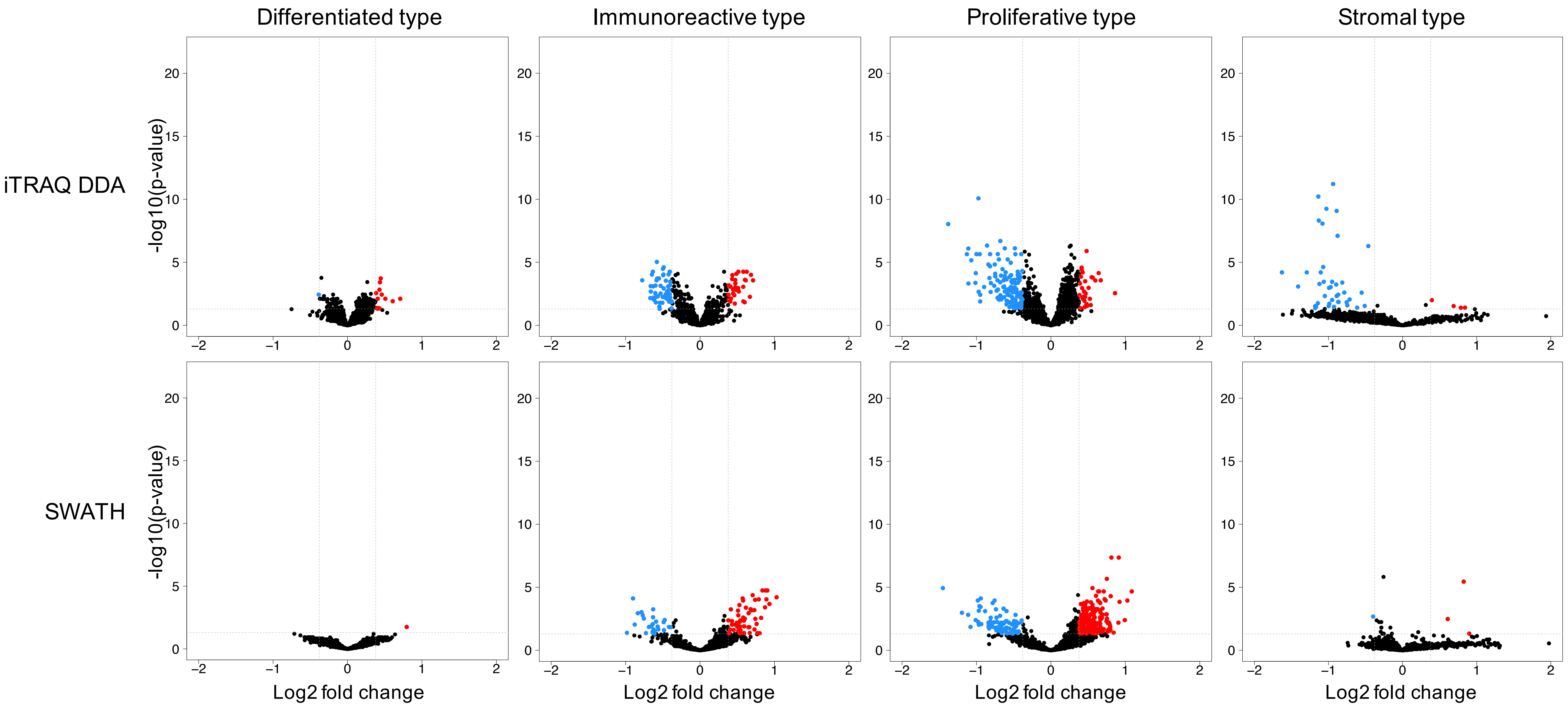
